# The ACE-inhibitor drug captopril inhibits ACN-1 to control dauer formation and aging

**DOI:** 10.1101/2023.07.17.549402

**Authors:** Brian M. Egan, Franziska Pohl, Xavier Anderson, Shoshana C. Williams, Imienreluefe Gregory Adodo, Patrick Hunt, Zuoxu Wang, Chen-Hao Chiu, Andrea Scharf, Matthew Mosley, Sandeep Kumar, Daniel L. Schneider, Hideji Fujiwara, Fong-Fu Hsu, Kerry Kornfeld

## Abstract

The renin-angiotensin-aldosterone system (RAAS) plays a well-characterized role regulating blood pressure in mammals. Pharmacological and genetic manipulation of the RAAS has been shown to extend lifespan in *C. elegans*, *Drosophila*, and rodents, but its mechanism is not well defined. Here we investigate the angiotensin-converting enzyme (ACE) inhibitor drug captopril, which extends lifespan in worms and mice. To investigate the mechanism, we performed a forward genetic screen for captopril hypersensitive mutants. We identified a missense mutation that causes a partial loss-of-function of the *daf-2* receptor tyrosine kinase gene, a powerful regulator of aging. The homologous mutation in the human insulin receptor causes Donohue syndrome, establishing these mutant worms as an invertebrate model of this disease. Captopril functions in *C. elegans* by inhibiting ACN-1, the worm homolog of ACE. Reducing the activity of *acn-1* via captopril or RNAi promoted dauer larvae formation, suggesting *acn-1* is a *daf* gene. Captopril-mediated lifespan extension xwas abrogated by *daf-16(lf)* and *daf-12(lf)* mutations. Our results indicate that captopril and *acn-1* control aging by modulating dauer formation pathways. We speculate that this represents a conserved mechanism of lifespan control.

**Summary Statement:** Captopril and *acn-1* control aging. By demonstrating they regulate dauer formation and interact with *daf* genes, including a new DAF-2(A261V) mutant corresponding to a human disease variant, we clarified the mechanism.

## Introduction

Aging is characterized by progressive degeneration of tissue structure and function that leads inexorably to death. A vital goal of aging research is to determine the mechanistic basis of age-related degeneration, since this information can help develop interventions that extend health span and lifespan. The renin-angiotensin-aldosterone system (RAAS) in mammals has been characterized extensively for its role in blood pressure regulation (Basso and Terragno, 2001; Igic, 2009). More recently, this pathway has been demonstrated to influence aging in a wide range of organisms; Egan *et al*. (2022) recently reviewed emerging evidence that this pathway plays a conserved role in longevity control in worms, flies, and rodents (Egan et al., 2022). However, the mechanism of longevity control has not been well defined in any animal, and this is now an important research objective.

Kumar *et al*. (2016) reported that captopril, an angiotensin-converting enzyme (ACE) inhibitor, extended *C. elegans* mean lifespan by ∼23% and maximum lifespan by ∼18% (Kumar et al., 2016). The first of what is now a large class of ACE inhibitors, captopril is an oligopeptide derivative developed in 1975 based on a peptide found in pit viper venom (Ondetti et al., 1977; Cushman and Ondetti, 1991). ACE inhibitors modulate the RAAS, a mechanism by which the body adapts to hypotension (Skeggs et al., 1956; Peach, 1977; Reid et al., 1978; Basso and Terragno, 2001; Fyhrquist and Saijonmaa, 2008; Igic, 2009; Zehnder et al., 2009). In response to a decline in blood pressure, the kidney releases renin, which cleaves angiotensinogen to angiotensin I. ACE converts angiotensin I to angiotensin II, and angiotensin II binds two transmembrane receptors – the primary effect is to stimulate aldosterone secretion and promote vasoconstriction to increase blood pressure. By blocking ACE and preventing the conversion of angiotensin I to angiotensin II, captopril lowers blood pressure (**Fig. 1A**).

**Figure 1:**
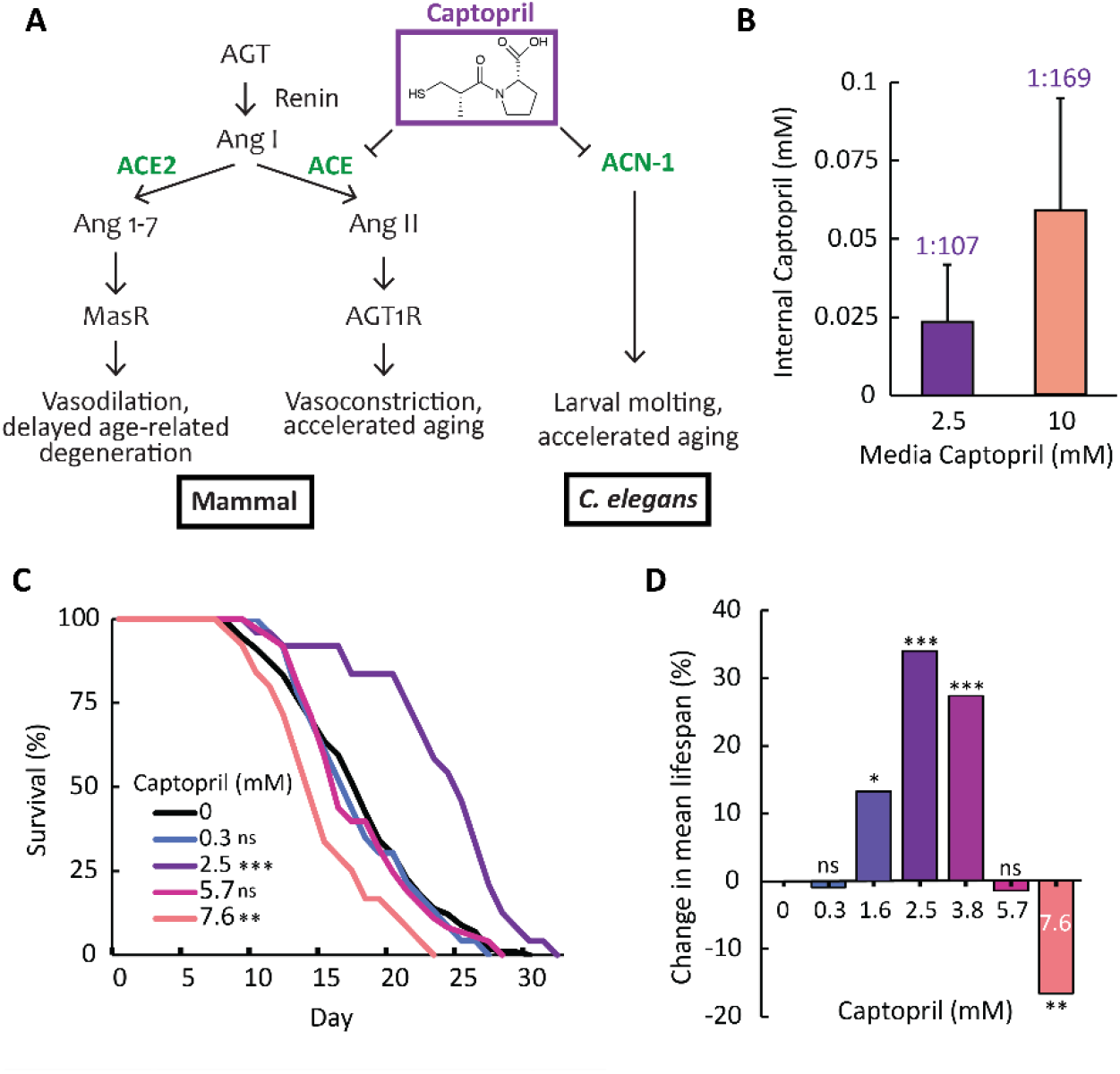
The ACE inhibitor captopril causes dose-dependent effects in *C. elegans*. **(A)** Captopril inhibits ACE (Angiotensin-Converting Enzyme) in mammals and ACN-1 (ACE-like Non-metallopeptidase) in *C. elegans*. AGT, angiotensinogen; Ang I, angiotensin I; ACE2, angiotensin-converting enzyme 2; Ang 1-7, angiotensin 1-7; Ang II, angiotensin II; MasR, Mas receptor; AGT1R, angiotensin II type-I receptor. **(B)** The internal concentration of captopril in *C. elegans* was measured by high-performance liquid chromatography-mass spectroscopy (HPLC-MS). Media Captopril indicates the concentration in the agar medium used to culture worms; Internal Captopril indicates the concentration in whole-animal lysates. Numbers are the ratio of internal captopril to media captopril, an indication of the ability of the animals to exclude captopril. Bars represent average and s.d. n=4 biological replicates, Student’s *t*-test. **(C,D)** Wild-type animals were cultured on NGM dishes seeded with *E. coli* bacteria and the indicated concentration of captopril in the medium. **(C)** Values indicate the fraction of the starting population that remains alive. **(D)** Bars represent the average change in mean lifespan compared to 0 mM control; positive values indicate lifespan extension, and negative values indicate lifespan reduction (see **Table S1** for statistics). ns, non-significant, p≥0.05; *, p<0.05; **, p<0.01; ***, p<0.001.

In Drosophila, drugs that inhibit the ACE pathway and mutations in ACE pathway genes can both extend lifespan (Ederer et al., 2018; Gabrawy et al., 2019, 2022). It has also been reported that the ACE inhibitor enalapril and the angiotensin II receptor antagonist losartan can extend the lifespan of normotensive mice and rats and delay age-related degeneration of tissue structure and function in the kidney, cardiovascular system, liver, and brain (Ferder et al., 1993, 1994; Inserra et al., 1995; González Bosc et al., 2000; Ferder et al., 2002; de Cavanagh et al., 2003; Inserra, 2003; Carter et al., 2004; Basso et al., 2005, 2007; de Cavanagh et al., 2008; Inserra et al., 2009; Santos et al., 2009; Carter et al., 2011; de Cavanagh et al., 2011; Keller et al., 2019). Based on findings in *C. elegans* (Kumar et al., 2016), the Intervention Testing Program (ITP) of the National Institute on Aging tested captopril in normotensive mice; captopril significantly increased the median and maximum lifespan of female and male mice (Strong et al., 2022). Genetic studies provide important support for these pharmacology studies, since disruption of the angiotensin II type I receptor (AGT_1_R) promotes longevity in normotensive mice (Benigni et al., 2009; Mattson and Maudsley, 2009; Nishiyama et al., 2009; Benigni et al., 2010; Cassis et al., 2010; Yabumoto et al., 2015). These results may be relevant to humans, since polymorphisms in the angiotensin II type I receptor gene are associated with extreme human longevity (Rigat et al., 1990; Benigni et al., 2013; Garatachea et al., 2013; Revelas et al., 2018).

Isaac and colleagues reported in 2003 that *C. elegans acn-1* (ACE-like non-metallopeptidase) encodes the homolog of mammalian ACE (Brooks et al., 2003). To analyze the function of *acn-1*, Isaac and colleagues used *acn-1* RNAi (Brooks et al., 2003): larvae arrested at the L2 stage with unshed L1 cuticle, indicating a defect in molting. Delivering *acn-1* RNAi at later stages revealed cuticle defects in L3/L4 larvae and adults, indicating *acn-1* functions to promote molting at multiple stages of larval development. Furthermore, seam cells displayed fusion defects, indicating *acn-1* plays a role in establishing cell fates. Consistent with this analysis of molting, Ruvkun and colleagues identified *acn-1* in a genome wide RNAi screen for molting defects (Frand et al., 2005). Analysis of ACN-1 transcriptional and translation reporters revealed that ACN-1 is expressed in hypodermal seam and excretory gland cells in embryos and larvae, as well as in the developing vulva and male tail (Brooks et al., 2003; Frand et al., 2005). Slack and colleagues reported in 2017 that *acn-1* interacts with heterochronic pathways (Metheetrairut et al., 2017). *acn-1* RNAi suppresses *let-7(lf)* lethality and seam cell defects and enhances the precocious phenotype of *hbl-1(lf)*. Loss of *apl-1* causes a similar phenotype, and loss of both *apl-1* and *acn-1* caused additive or synergistic defects. Reducing *acn-1* activity reduces expression of *apl-1*, suggesting *acn-1* functions upstream to control *apl-1* expression. These results are consistent with two reports of global analysis that identified *acn-1* among a set of genes with cyclic larval expression (Kim et al., 2013; Hendriks et al., 2014). These studies led to the model that *acn-1* is a heterochronic gene that promotes larval seam cell fates by functioning downstream of *let-7* and upstream of *apl-1*.

To test the hypothesis that *acn-1* regulates lifespan, Kumar *et al*. (2016) used RNAi to reduce *acn-1* activity (Kumar et al., 2016). Wild-type animals cultured with *acn-1* RNAi starting at the embryonic stage displayed significantly extended lifespan (Kumar et al., 2016). Furthermore, *acn-1* functions in adults to extend lifespan, since animals cultured with RNAi bacteria starting at the L4 stage also display an extended lifespan; this indicates that, in addition to its regulation of molting and development in larvae, *acn-1* continues to be expressed in adults to control aging. *acn-1* RNAi delayed age-related declines of body movement and pharyngeal pumping and increased resistance to heat and oxidative stress, indicating *acn-1* activity accelerates aging and reduces stress resistance (Kumar et al., 2016). If captopril treatment and *acn-1* RNAi extend life span by a similar mechanism, then combining treatments is predicted to be non-additive. As predicted, captopril did not further extend the life span of animals treated with *acn-1* RNAi (Kumar et al., 2016). Based on these findings, Kumar *et al*. (2016) hypothesized that *acn-1* controls lifespan, and captopril extends lifespan by inhibiting ACN-1 activity (**Fig. 1A**) (Kumar et al., 2016). Captopril and other ACE inhibitors have been the topic of intense study in recent years as one of its targets, ACE2, is the cellular point-of-entry for the SARS-CoV-2 coronavirus responsible for the COVID-19 pandemic (Wrapp et al., 2020).

To investigate the mechanism of captopril in longevity control, we performed dose-response experiments and exploited this information to design a forward genetic screen for mutants that are hypersensitive to captopril. We identified *am326*, a novel missense allele of the *daf-2* gene, which encodes the *C. elegans* homolog of the insulin/IGF-1 receptor. The *daf-2(am326)* mutation results in an alanine to valine substitution at amino acid 261, a residue in the extracellular L1 ligand binding domain of the DAF-2 protein. This alanine residue is conserved in the human insulin receptor (INSR) at amino acid position 119 (or position 92 following the termination of the signal sequence), and human patients with the identical alanine to valine substitution suffer from Donohue syndrome (also known as “leprechaunism”), a disease characterized by growth restriction, morphological abnormalities, reduction of glucose homeostasis, and early death (<1 year of life) (Donohue, 1948; Longo et al., 2002). *daf-2* mutations cause an extended lifespan and a dauer constitutive (Daf-c) phenotype; mutant animals enter the dauer stage under conditions where wild type animals undergo reproductive development (Cassada and Russell, 1975; Klass and Hirsh, 1976; Klass, 1977; Kenyon et al., 1993; Kimura et al., 1997). Our results indicate that captopril treatment and reducing the activity of *acn-1* by RNAi also cause a Daf-c phenotype, indicating that *acn-1* controls both dauer formation and adult longevity. The lifespan effects of captopril require the *daf-16* and *daf-12* genes, providing further evidence that *acn-1* interacts with the dauer pathway to promote longevity. Together, our results advance the understanding of the mechanism of action of the ACE pathway in longevity control.

## Results

### High dose captopril causes developmental arrest

Captopril is an established regulator of aging, since treatment can extend the lifespan of *C. elegans* and mice (Kumar et al., 2016; Egan et al., 2022; Strong et al., 2022). The optimal dose for extending longevity in *C. elegans*, 2.5 mM dissolved in nematode growth-medium (NGM), increased lifespan by more than 30% (**Fig. 1C,D, Table S1**) and delayed the age-related decline of pharyngeal pumping and body movement (Kumar et al., 2016). This dose also increases heat and oxidative stress resistance (Kumar et al., 2016). The longevity effect is dose-dependent; reducing the concentration to 0.3 mM or increasing the concentration to 5.7 mM abrogated the lifespan extension (**Fig. 1C,D, Table S1**). At even higher doses, captopril delayed larval development, reduced length of animals at adulthood, and reduced lifespan compared to untreated animals, indicating dose dependent toxicity (**Fig. 1C,D, S1A,B, Table S1**). When cultured on 16 mM captopril, most wild-type worms grew to adulthood and produced progeny by day 3 post-hatch (**Fig. 2B, S1B**). However, at a higher dose of 19 mM captopril, wild-type worms displayed high rates of lethality and were unable to reach adulthood (**Fig. 2B**).

**Figure 2:**
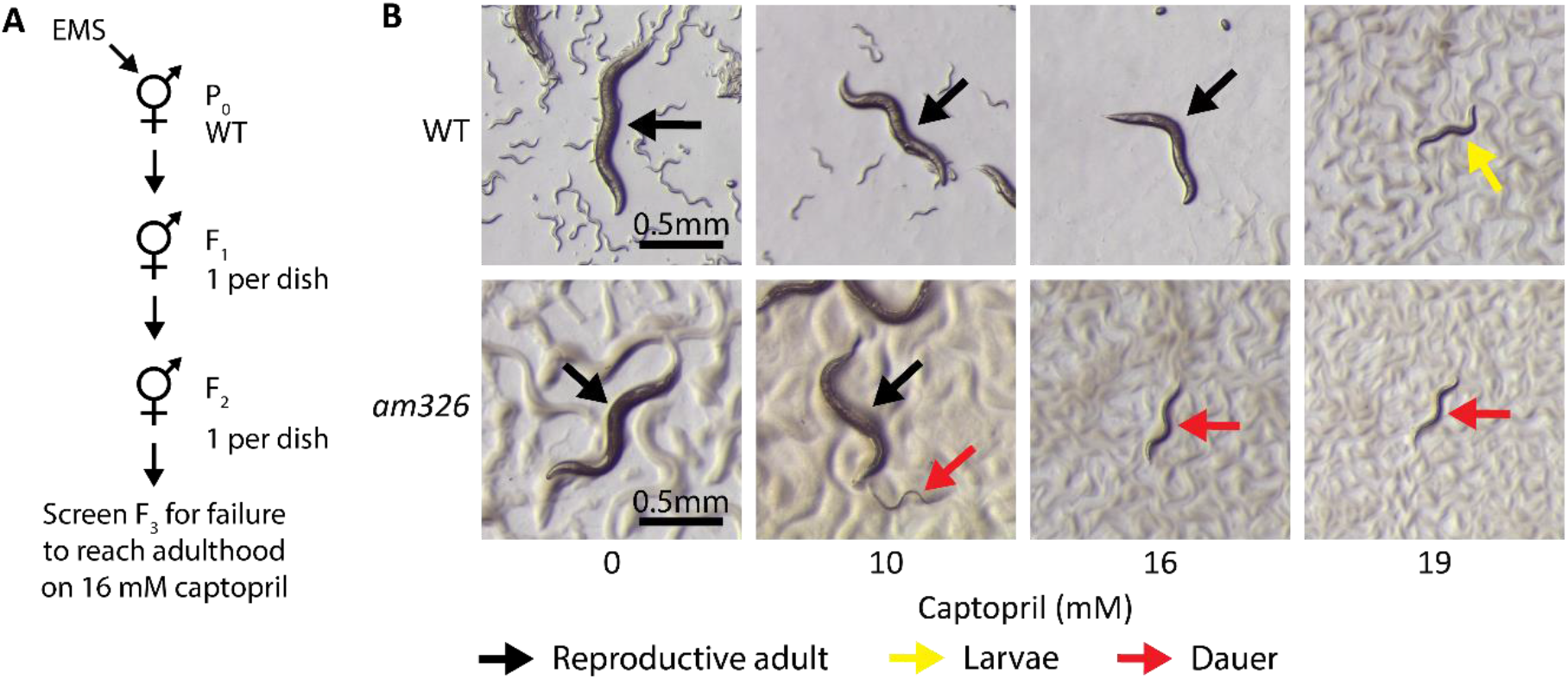
A forward genetic screen for mutants hypersensitive to high dose captopril. **(A)** Diagram of a forward genetic screen for mutant animals that are hypersensitive to high-dose captopril. Wild-type L4 hermaphrodites were mutagenized with ethyl methanesulfonate (EMS), and clonal populations derived from individual F2 self-progeny were analyzed for failure to grow to adulthood on 16 mM captopril. **(B)** Brightfield images show WT and *am326* mutant animals cultured with the indicated concentration of captopril in the medium. WT animals mature to adulthood and produce progeny when cultured with 0, 10, and 16 mM captopril (black arrows) but display larval arrest when cultured with 19 mM captopril (yellow arrow). *am326* mutant animals mature to adulthood and produce progeny when cultured with 0 and 10 mM captopril (black arrows), but develop as dauer larvae when cultured with 16 and 19 mM captopril (red arrow). Some *am326* animals enter dauer when cultured with 10 mM captopril. Scale bar=0.5 mm.

### Determination of internal captopril concentration

*C. elegans* dwells in a chemically complex soil environment and is adept at excluding xenobiotic compounds. Thus, the concentration of a drug inside the worms is typically much lower than the concentration in its environment, often 2-4 orders of magnitude lower (Lindblom and Dodd, 2006; Burns et al., 2010; Xiong et al., 2017). The concentration of captopril in the medium that is necessary for lifespan extension is well characterized, but the concentration inside the worms is unknown. To determine this value, we cultured a large population of animals with 2.5 and 10 mM captopril in the medium, collected a whole-worm lysate, and used High Performance Liquid Chromatography/Mass Spectroscopy (HPLC-MS) to measure the internal concentration of captopril. When the external concentration was 2.5 mM, the concentration in worm lysate was ∼0.02 mM; this value is 107-fold lower than the external concentration, demonstrating that these animals are effective at excluding captopril (**Fig. 1B**). This value is less than one order of magnitude higher than the serum concentration in humans treated with captopril (∼0.002-0.003 mM), indicating that the lifespan extending concentration in nematodes is similar to the therapeutic concentration in humans (Bahmaei et al., 1997; Huang et al., 2006). Increasing the external concentration to 10 mM resulted in an internal concentration of ∼0.06 mM, 169-fold lower than the external concentration in the medium. Together the results indicate that the internal concentration of captopril is dose-dependent (**Fig. 1B**).

### A forward-genetic screen for captopril-hypersensitive mutants

To investigate the mechanism of lifespan extension caused by captopril, we performed a genetic screen for mutants with an abnormal dose response to the drug. The aging-related phenotypes caused by captopril, such as increased lifespan or delayed decline of body movement rate, are laborious to score and not suitable for such a genetic screen. Instead, we chose to screen for mutations that caused hypersensitivity to captopril larval arrest/lethality, since this phenotype can be readily scored. 16 mM captopril in the medium is the highest dose at which the majority of wild-type animals survive to adulthood and produce progeny (**Fig. 2B, S1B**); thus, we performed a clonal screen for populations of mutant animals that failed to survive and reproduce at this dose (**Fig. 2A**).

Wild-type L4 hermaphrodites were exposed to a chemical mutagen, 50 mM ethylmethansulfanate (EMS) as described in (Brenner, 1974). These P_0_ animals were allowed to recover and reproduce, and their F_1_ self-progeny were moved to individual dishes. F_2_ progeny were moved to individual dishes and cultured until their F_3_ progeny reached adulthood. Dishes containing large amounts of F_3_ adults were treated with alkaline hypochlorite solution to yield eggs, and populations of F_4_ eggs were cultured on dishes containing 16 mM captopril. After 4-5 days, these dishes were evaluated using a dissecting microscope for populations that lacked fertile adults.

To control for mutants that fail to thrive under non-specific stress conditions, we incorporated control dishes with a metal stress environment into the genetic screen. The same F4 population that was challenged with 16 mM captopril was also cultured on dishes containing high-dose manganese, high-dose zinc, or zinc deficiency caused by a zinc chelator (TPEN). If a mutant strain failed to reproduce in more than one stress condition, the mutant was discarded. After screening ∼1,950 haploid mutant genomes, we identified one mutant strain that displayed specific hypersensitivity to captopril; we designated this mutation *am326* (**Fig. 2B**). This strain was backcrossed three times to the wild-type N2 strain, and the resulting strain (WU1939) was used for all future experiments.

### *am326*, a mutation that causes hypersensitivity to captopril, also causes lifespan extension and promotes dauer development

When cultured with high-dose captopril, *am326* mutant animals failed to mature into adults and produced no progeny after 5 days (**Fig. 2B**). We noticed that these mutant animals formed dauer larvae, as determined by morphology observed with a dissecting microscope and survival following treatment with sodium dodecyl sulfate (SDS) (Cassada and Russell, 1975; Riddle et al., 1981; Swanson and Riddle, 1981; Vowels and Thomas, 1992; Hu, 2018). Dauer is an alternative L3 larval stage that *C. elegans* can enter when conditions are not optimal for reproductive development (Cassada and Russell, 1975; Klass and Hirsh, 1976; Klass, 1977). These conditions include high population density, low resource availability, and elevated temperature (Cassada and Russell, 1975; Klass and Hirsh, 1976; Klass, 1977). Dauer animals are stress resistant and extremely long-lived (Cassada and Russell, 1975; Klass and Hirsh, 1976; Klass, 1977; Larsen, 1993). *am326* mutants displayed high rates of dauer formation when cultured with 16 mM captopril; by contrast, wild-type animals did not form dauer larvae under these conditions (**Fig. 2B, 3A, Table S2**). Dauer formation was dose dependent; the percentage of dauer animals in a population increased as the captopril concentration increased (**Fig. 3A, Table S2**).

**Figure 3:**
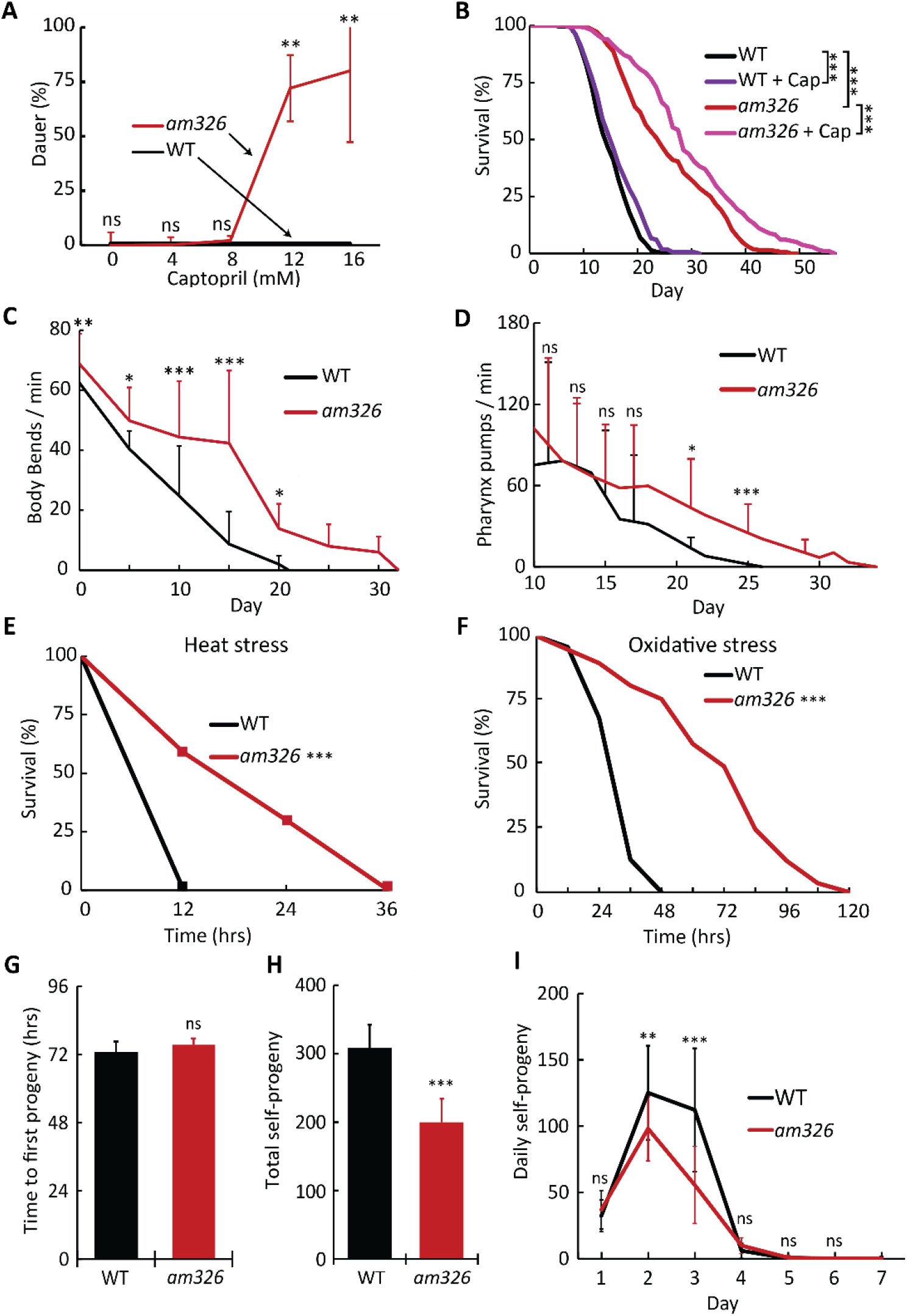
Analysis of *am326* phenotypes. **(A)** WT (black line) and *am326* mutant (red line) embryos were cultured on dishes containing the indicated concentration of captopril at 20°C. Values are the average percentage of animals in the dauer stage and s.d. (see **Table S2** for statistics). **(B)** WT and *am326* mutant animals were cultured with 0 or 2.5 mM captopril (+ Cap). Values indicate the fraction of the starting population that remains alive (see **Table S3** for statistics). **(C)** Movement of WT and *am326* mutant animals was measured by counting body turns in 15 s intervals using a dissecting microscope throughout adulthood. Values are body bends per minute and s.d. n=25-40 animals, one-way ANOVA with Tukey’s post-hoc HSD. **(D)** Pharyngeal pumping rate of WT and *am326* mutant animals was measured in 15 s intervals using a dissecting microscope throughout adulthood. Values are average pharynx pumps per minute and s.d. n=27-30 animals, one-way ANOVA with Tukey’s post-hoc HSD. **(E)** Thermotolerance was measured by culturing WT and *am326* mutant embryos at 20°C for 72 hrs (until ∼adult day 0), shifting animals to 35°C, and measuring survival every 12 hrs. Values indicate the fraction of the starting population that remains alive (see **Table S3** for statistics). **(F)** Resistance to oxidative stress was measured by culturing WT and *am326* embryos on standard medium for 72 hrs (until ∼adult day 0), transferring animals to medium containing 40 mM paraquat, and measuring survival every 12 hrs. In this experiment, animals that died from matricidal hatching were not censored because matricidal hatching was the primary cause of death for paraquat-treated animals. Values indicate the fraction of the starting population that remains alive (see **Table S3** for statistics). **(G)** Freshly laid eggs from WT and *am326* hermaphrodites were cultured for 65 hrs, then the resulting self-fertile adults were observed hourly to determine the time when they laid their first egg. Values are average time to first egg lay and s.d. n=15-16 animals, Student’s *t-*test. **(H,I)** Total brood size and daily progeny production of self-fertile hermaphrodites were measured by counting the number of viable self-progeny produced by WT and *am326* hermaphrodites throughout their reproductive period. Values are average number of progeny and s.d. n=14 animals. H, Student’s *t-*test; I, one-way ANOVA with Tukey’s post-hoc HSD. ns, non-significant, p≥0.05; *, p<0.05; **, p<0.01; ***, p<0.001.

To investigate the role of the *am326* mutation in aging, we measured age-related phenotypes. Homozygous *am326* mutant animals displayed an increased lifespan of 74.5% compared to wild type; captopril treatment further extends the lifespan of *am326* mutants, indicating the mutation does not abrogate the effect of the drug (**Fig. 3B, Table S3**). To analyze health span, we measured the age-related declines of body movement and pharynx pumping. Homozygous *am326* mutant animals displayed reduced age-related declines in body movement and pharyngeal pumping; these phenotypes are only apparent after mid-life (**Fig. 3C,D**). Some mutations that extend lifespan also increase stress resistance (Lithgow et al., 1995; Johnson et al., 2000). Homozygous *am326* mutant animals displayed increased heat and oxidative stress resistance (**Fig. 3E,F, Table S3**). Some mutations that extend lifespan also delay development, reduce fecundity, and/or extend the reproductive period (Huang et al., 2004; Hughes et al., 2011; Luo and Murphy, 2011). Homozygous *am326* mutants did not display delayed development compared to wild type, as determined by measuring the duration from hatching to egg-laying (**Fig. 3G**). Homozygous *am326* mutant hermaphrodites did display a 35% reduction in self-progeny number compared to wild type (**Fig. 3H**); however, these mutants did not display an extension of the reproductive period (**Fig. 3I**). Thus, the *am326* mutation causes a large magnitude lifespan extension, extension of neuromuscular health span, increased stress resistance, and a moderate reduction in brood size, but does not cause a delay in developmental or reproduction.

### *am326* is a previously uncharacterized missense mutation in the *daf-2* receptor-tyrosine kinase gene

To identify the molecular lesion in the *am326* strain, we utilized an EMS density mapping strategy (Zuryn et al., 2010). We performed multiple independent backcrosses to the wild-type N2 strain using the temperature-dependent dauer formation phenotype to homozygose *am326/am326* animals. Using whole genome sequencing, we generated a list of 45 base pair variants that were present in all backcrossed *am326/am326* strains but were absent from wild type, indicating they were caused by the EMS mutagenesis and are strongly linked to *am326* (**Table S4**). 36 variants were clustered in a region of chromosome III between 2.5 and 5.6 Mbp, suggesting that one of the variants in this region was likely the causative mutation. The remaining 9 variants were dispersed on other chromosomes (**Fig. 4A, Table S4**). We predicted the effect of each variant on gene activity; variants outside coding regions (introns, 5’ untranslated regions, intergenic regions, or non-coding RNA) were excluded. Eight variants affected exons, including one missense change in the *daf-2* gene. This mutation is a single nucleotide change (cytosine to thymine) at position 782, resulting in an alanine (GCG) to valine (GTG) substitution. The lesion occurs in exon 7, which encodes the extracellular L1 ligand binding domain of the DAF-2 protein (**Fig. 4C**).

**Figure 4:**
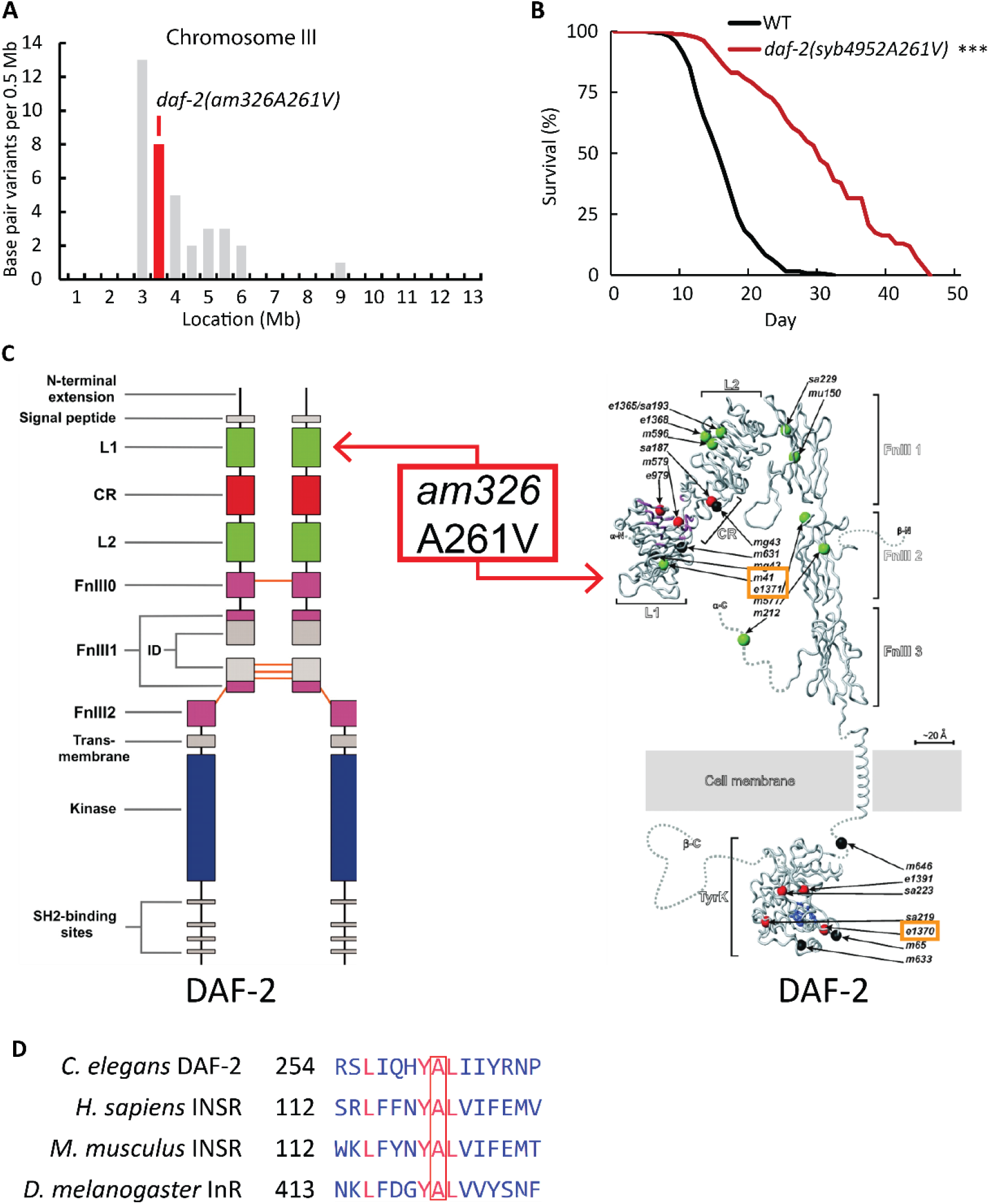
*am326* is an allele of *daf-2*. **(A)** The x-axis represents the location on chromosome III in megabases (Mb). Bars indicate the number of base pair variants in the *am326* strain in 500,000 base pair bins. Red indicates the location of *am326*, an allele of *daf-2*, at position 3,015,386 bp (see **Table S4** for details). **(B)** WT and *daf-2(syb4952A261V)*, a CRISPR-generated mutant strain, were analyzed for survival. Values indicate the fraction of the starting population that remains alive. (***, p<0.001; see **Table S5** for statistics). **(C)** Left, a diagram of the predicted DAF-2 protein as a dimer with domains colored and labelled. The *am326* mutation causes an alanine to valine substitution at amino acid 261 that affects the L1 extracellular ligand-binding domain (L1BD); right, a structure of the DAF-2 protein showing the location of the alanine to valine substitution in the L1BD and the location of other amino acid substitutions caused by previously characterized mutations of *daf-2*. Orange boxes: *daf-2* mutations used in this study. Adapted from (Patel et al., 2008). **(D)** Multiple sequence alignment of DAF-2 and homologs in humans, mice, and *Drosophila*. Conserved amino acids are red, and Alanine 261 is marked by the box. Alignment was performed using COBALT (Papadopoulos and Agarwala, 2007).

DAF-2 is a well-studied member of the insulin/IGF-1 signaling (IIS) pathway; numerous partial loss-of-function mutations in this gene have been shown to affect aging, development, and dauer entry (Kenyon et al., 1993; Gems et al., 1998; Patel et al., 2008). We hypothesized that the missense mutation in *daf-2* is the cause of the *am326* mutant phenotype, since *daf-2* mutant animals have been previously demonstrated to cause phenotypes including Daf-c, extended lifespan, stress resistance, and reduced brood size (Kenyon et al., 1993; Dorman et al., 1995; Kimura et al., 1997) (reviewed in (Gems et al., 1998; Patel et al., 2008)). To test this hypothesis, we performed complementation assays. Male P_0_ *am326* animals were crossed to hermaphrodite P_0_ animals with *daf-2(lf)* mutations, as well as hermaphrodite P_0_ animals with an *age-1(lf)* mutation as a control; *age-1(lf)* mutants also display temperature-dependent dauer formation (Friedman and Johnson, 1988a, 1988b; Klass, 1983). Heterozygous F_1_ larvae were temperature-shifted to 25°C for 48 hours and observed for dauer formation. **Table 1** shows that *am326* failed to complement *daf-2(lf)* mutations, but complemented an *age-1(lf)* mutation, indicating that the *am326* mutation is an allele of the *daf-2* gene.

**Table 1:**
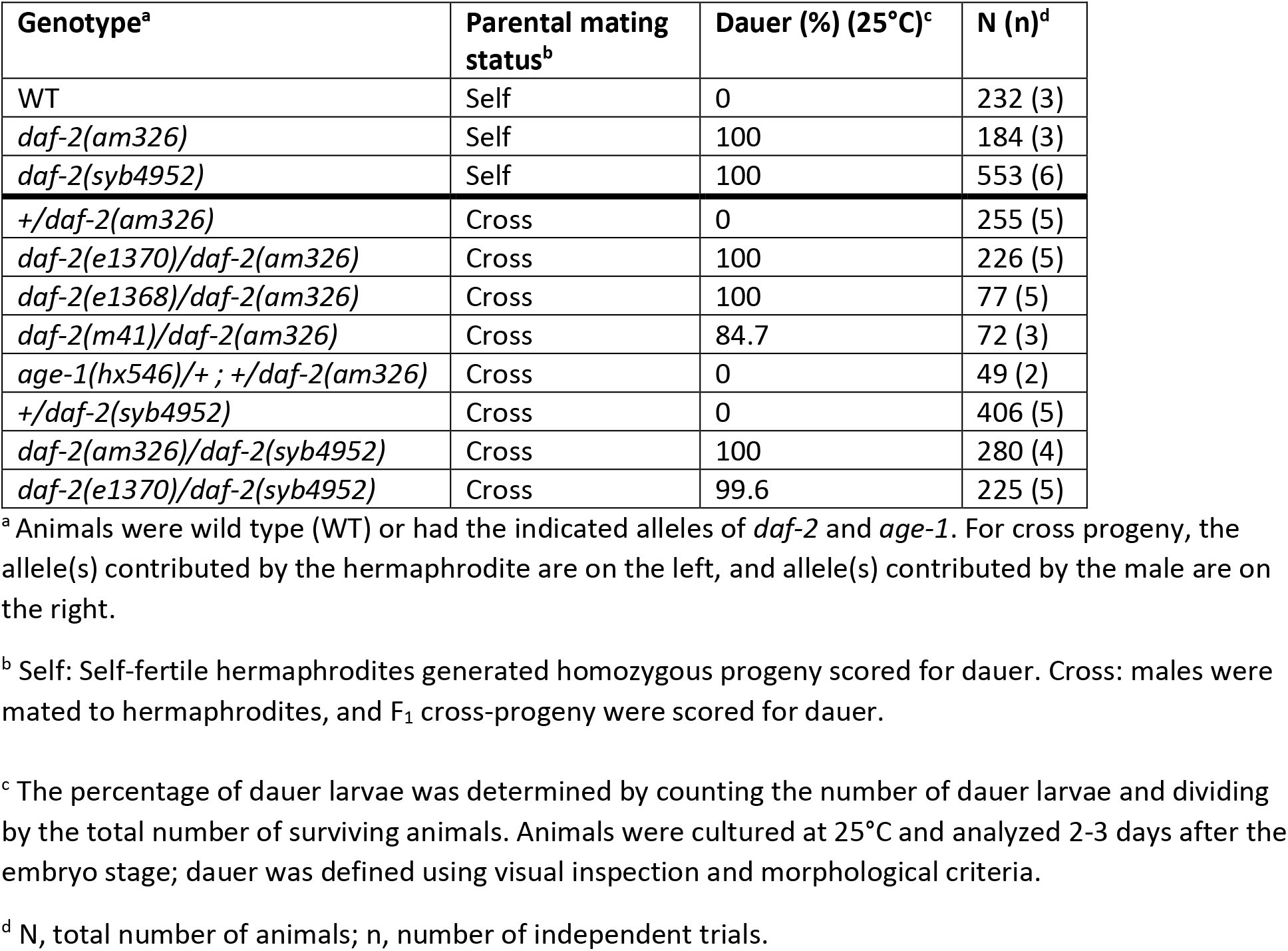
Complementation tests indicate that *am326* is an allele of *daf-2*

| Genotype<sup>a</sup> | Parental mating status<sup>b</sup> | Dauer (%) (25°C)<sup>c</sup> | N (n)<sup>d</sup> |
| --- | --- | --- | --- |
| WT | Self | 0 | 232 (3) |
| *daf-2(am326)* | Self | 100 | 184 (3) |
| *daf-2(syb4952)* | Self | 100 | 553 (6) |
| *+/daf-2(am326)* | Cross | 0 | 255 (5) |
| *daf-2(e1370)/daf-2(am326)* | Cross | 100 | 226 (5) |
| *daf-2(e1368)/daf-2(am326)* | Cross | 100 | 77 (5) |
| *daf-2(m41)/daf-2(am326)* | Cross | 84.7 | 72 (3) |
| *age-1(hx546)/+ ; +/daf-2(am326)* | Cross | 0 | 49 (2) |
| *+/daf-2(syb4952)* | Cross | 0 | 406 (5) |
| *daf-2(am326)/daf-2(syb4952)* | Cross | 100 | 280 (4) |
| *daf-2(e1370)/daf-2(syb4952)* | Cross | 99.6 | 225 (5) |

<sup>a</sup> Animals were wild type (WT) or had the indicated alleles of *daf-2* and *age-1*. For cross progeny, the allele(s) contributed by the hermaphrodite are on the left, and allele(s) contributed by the male are on the right.
<sup>b</sup> Self: Self-fertile hermaphrodites generated homozygous progeny scored for dauer. Cross: males were mated to hermaphrodites, and F<sub>1</sub> cross-progeny were scored for dauer.
<sup>c</sup> The percentage of dauer larvae was determined by counting the number of dauer larvae and dividing by the total number of surviving animals. Animals were cultured at 25°C and analyzed 2-3 days after the embryo stage; dauer was defined using visual inspection and morphological criteria.
<sup>d</sup> N, total number of animals; n, number of independent trials.

DNA sequencing was used to confirm the presence of the *daf-2(am326A261V)* allele in the originally isolated *am326* mutant strain and in four serially backcrossed mutant strains (WU1975-WU1978); this allele was not present in the wild-type strain used for EMS mutagenesis, indicating a strong correlation between the *daf-2(am326A261V)* allele and the Daf-c phenotype. Because alanine and valine are similar amino acids, we wanted to rigorously confirm that this missense change causes the phenotype. CRISPR genome editing was used to insert this base change into the endogenous *daf-2* locus of a wild-type N2 strain. The resulting *daf-2(syb4952A261V)* mutant animals displayed a 100% penetrant Daf-c phenotype (**Table 1**). Furthermore, the *daf-2(syb4952)* allele failed to complement the *daf-2(am326)* and *daf-2(e1370)* mutations for the Daf-c phenotype, demonstrating that the A261V change caused a *daf-2* loss-of-function phenotype (**Table 1**). As expected, the *daf-2(syb4952)* mutant animals displayed temperature- and captopril-dependent dauer formation (**Table 1, S2**), and were long lived (**Fig. 4B, Table S5**). DAF-2 is the nematode homolog of the mammalian insulin receptor (INSR); we performed amino acid sequence alignment to examine if this amino acid was conserved in homologous proteins. Alanine 261, as well as the adjacent amino acids, are present in the homologous position in human and mouse INSR and the *Drosophila* insulin receptor (InR), indicating that this region is highly conserved (**Fig. 4D**).

### Captopril treatment influences dauer development

Some *daf-2* loss-of function mutations, such as *daf-2(e1370),* cause a temperature sensitive Daf-c phenotype (Kenyon et al., 1993; Patel et al., 2008). To evaluate the temperature sensitivity of the *daf-2(am326)* phenotype, we cultured animals at 20, 22, and 25°C and measured the percentage of dauer larvae in the population. The percentage of dauer larvae increased significantly from 0% at 20°C to 16.8% at 22°C and 90% at 25°C (**Fig. 5A, Table S2**). To investigate the interaction with captopril treatment, we cultured animals with 0, 4, 8, 12 or 16 mM captopril at the three temperatures. At 20 and 22°C, captopril treatment caused a dose-dependent increase in dauer formation (**Fig. 5A, Table S2**). At 25°C, the lowest dose of captopril (4 mM) resulted in constitutive dauer formation; at 16 mM captopril, the percentage of animals that entered dauer was reduced as a result of some animals undergoing larval arrest prior to the L2 stage. Thus, the dauer formation promoting effect of captopril is dose dependent and additive with high temperature.

**Figure 5:**
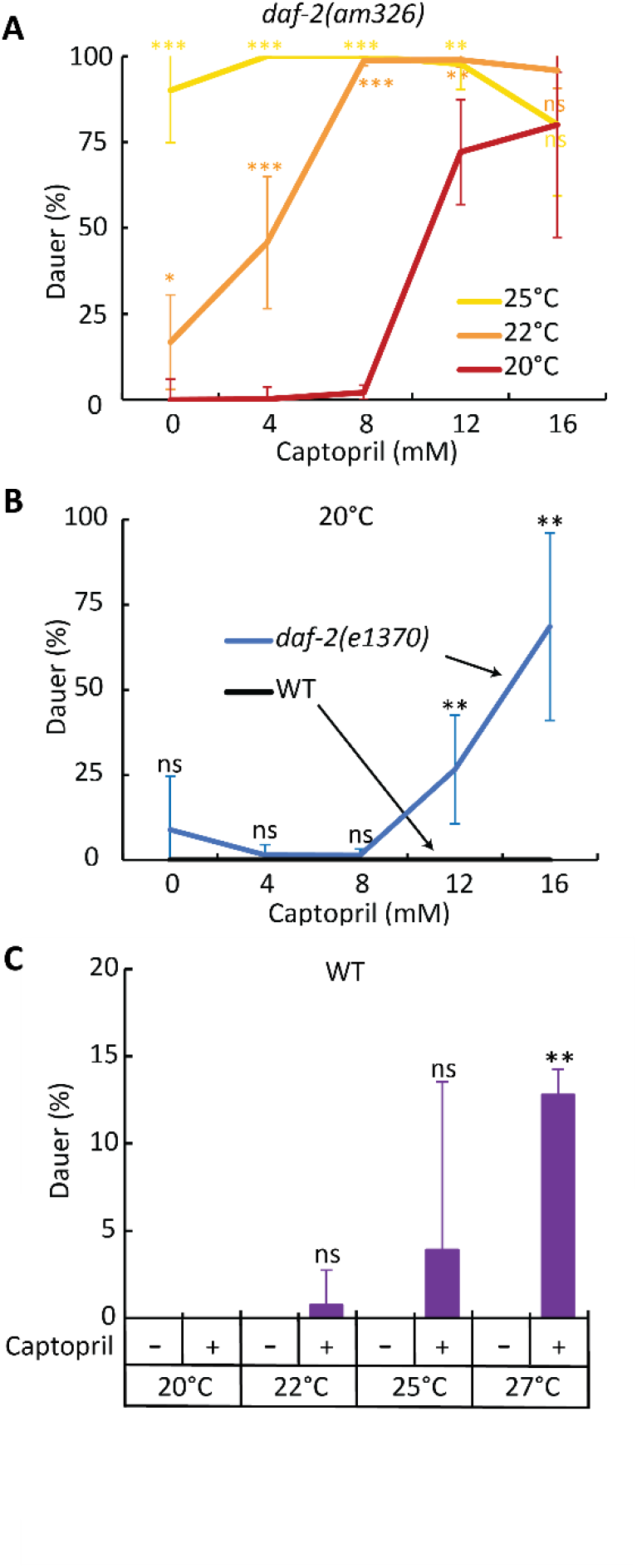
Captopril promotes dauer formation. **(A)***daf-2(am326)* embryos were cultured with the indicated concentration of captopril at 20°C (red), 22°C (orange) or 25°C (yellow). Values are the average percentage of animals in the dauer stage and s.d. At 22 and 25°C, animals treated with 16 mM captopril displayed less than 100% dauer because some animals arrested at developmental stages prior to L2. **(B)** WT (black line) and *daf-2(e1370)* mutant (blue line) embryos were cultured on dishes containing the indicated concentration of captopril at 20°C. Values are the average percentage of animals in the dauer stage and s.d. **(C)** WT embryos were cultured with either 0 (-) or 16 (+) mM captopril at the indicated temperatures. Values are the average percentage of animals in the dauer stage and s.d. (see **Table S2** for statistics on this figure). ns, non-significant, p≥0.05; *, p<0.05; **, p<0.01; ***, p<0.001.

To determine whether captopril-induced dauer formation was a specific effect of the *am326* allele or a general property of *daf-2(lf)* mutants, we measured the effect of captopril on *daf-2(e1370)*. Captopril induced dauer formation in a dose-dependent manner, suggesting that this effect is not specific to *daf-2(am326)* (**Fig. 5B, Table S2**).

To evaluate the specificity of the captopril effect on dauer formation, we analyzed a different small molecule drug that extends lifespan and is toxic at high doses, the anticonvulsant ethosuximide (Evason et al., 2005; Collins et al., 2008). Ethosuximide is an FDA-approved medication that was previously shown to extend lifespan in *C. elegans* at a similar concentration to captopril (2-4 mM). We exposed *daf-2(am326)* mutants to ethosuximide at concentrations up to 32 mM; no significant dose dependent effect on dauer formation was observed at 20, 22, or 25°C (**Fig. S2A,B, Table S6**). These results suggest the dauer formation phenotype is a distinctive effect of captopril and not a general xenobiotic stress response.

To determine if the dauer promoting effects of captopril require a *daf-2* mutation or can also be observed in wild-type animals, we used high temperature to sensitive wild-type animals to dauer formation and examined the effect of captopril. We cultured wild-type animals at 20, 22, 25 and 27°C, since 27°C was previously shown to induce dauer in HID (high-temperature dauer induction) mutants (Ailion and Thomas, 2003). At 22 and 25°C, 16 mM captopril treatment resulted in a small increase of dauer formation to 0.8% and 3.9%, respectively, that was not statistically significant with this sample size. At 27°C, 16 mM captopril significantly increased dauer formation to 12.8% (**Fig. 5C, Table S2**). Thus, the dauer promoting effect of captopril can be observed if animals are sensitized using a *daf-2* mutation or simply using high temperature in wild-type animals.

### The *acn-1* gene promotes dauer formation

Based on our model that captopril functions by inhibiting the *acn-1* gene (**Fig. 1A**), we hypothesized that reducing the activity of *acn-1* would promote dauer formation. To test this hypothesis, we treated *daf-2(am326)* mutants with bacteria expressing *acn-1* dsRNA and measured dauer formation. *acn-1* RNAi failed to induce the dauer state in wild-type animals at either 20, 22, or 25°C (**Fig 6A, Table S7**); this result is consistent with our observation that captopril fails to significantly induce dauer formation under similar conditions. However, *acn-1* RNAi significantly increased the percentage of dauer stage animals in *am326* mutants from 13.8% to 31.5% at 22°C (**Fig. 6A, Table S7**). This finding suggests that reducing *acn-1* activity through RNAi or inhibition with captopril promotes dauer formation under the sensitized conditions created by the *daf-2(am326)* mutation.

**Figure 6:**
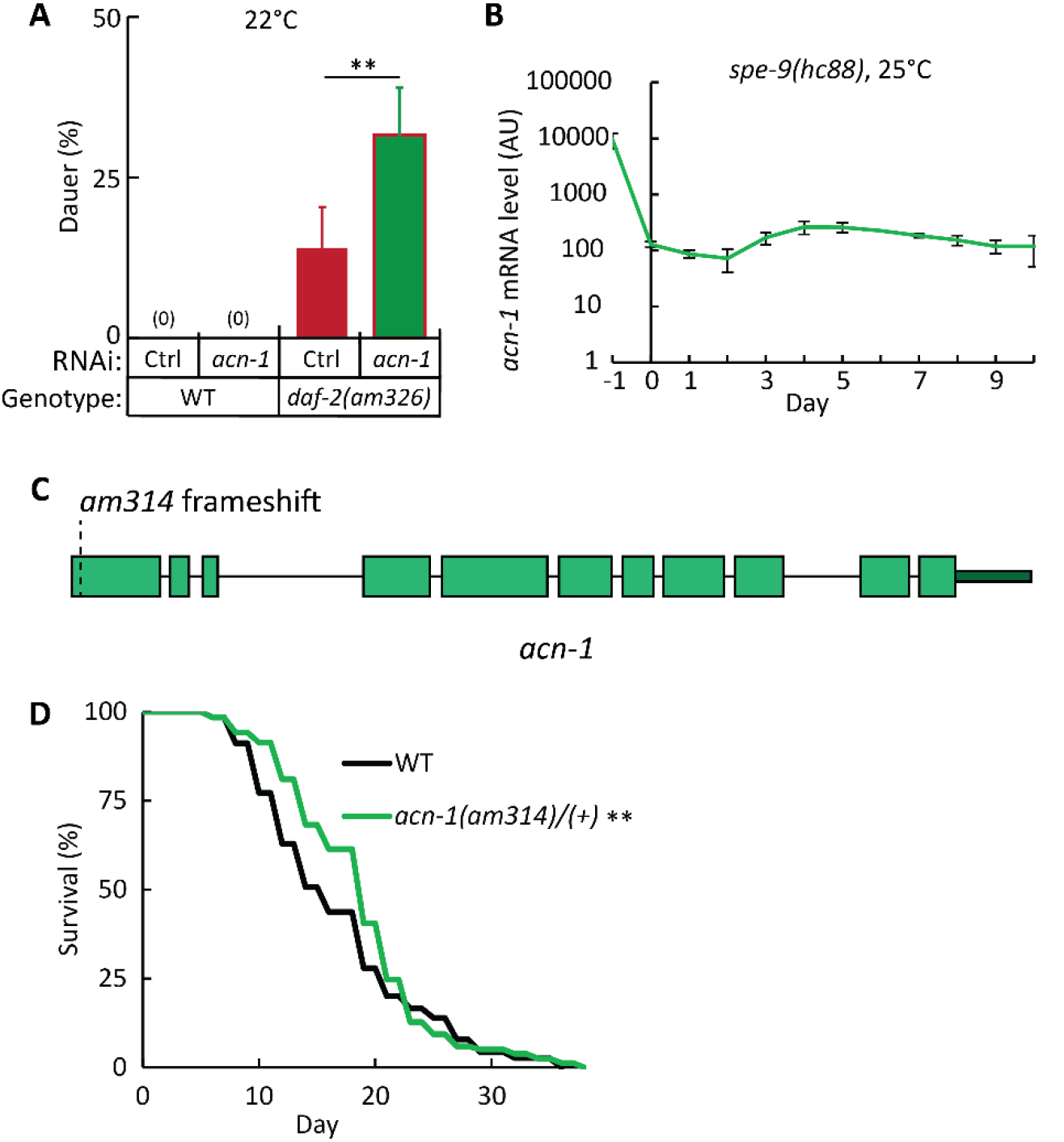
*acn-1* is expressed in larvae and adults and influences dauer formation and adult lifespan. **(A)** WT and *daf-2(am326)* embryos were cultured at 22°C with *E. coli* HT115 expressing either *acn-1* dsRNA or a vector control. Values are the average percentage of dauer larvae and s.d. (see **Table S7** for statistics). **(B)** *spe-9(hc88*) arrested at the L1 stage were transferred to dishes with abundant *E. coli* and cultured at 25°C until the indicated day; total RNA was analyzed by RNAseq. Values are *acn-1* mRNA level in arbitrary units (AU) and s.d. Days -1 and 0 were larval stages, and Day 1 was defined as the first day of adulthood; at 25°C, Day 1 occurs approximately two days after arrested L1 animals were transferred to food. **(C)** Diagram of the *acn-1* gene showing the location of the *am314* frameshift mutation. Green boxes, exons; black lines, introns; dark green box, 3’ UTR. **(D)** WT and animals heterozygous for *acn-1(am314)*, a CRISPR-generated mutation, were analyzed for survival. Values indicate the fraction of the starting population that remains alive (see **Table S5** for statistics). **, p<0.01.

To elucidate how reducing *acn-1* activity causes an extension of lifespan, we monitored *acn-1* expression daily beginning in larvae and extending until ∼day 12 of adulthood. *acn-1* mRNA levels were measured by RNA-seq in a synchronized population of *spe-9(lf)* animals raised at 25°C to prevent progeny production. *acn-1* mRNA levels were high in larvae, consistent with previous reports (**Fig. 6B**) (Brooks et al., 2003; Frand et al., 2005; Oskouian et al., 2005; Metheetrairut et al., 2017). Interestingly, *acn-1* mRNA levels declined in young adults but then remained relatively constant throughout adulthood (**Fig. 6B**). These results suggest that *acn-1* is expressed in young and old adults, consistent with an activity in adults that controls aging.

We previously used RNAi to investigate *acn-1* function in adults. To extend this analysis, we used CRISPR/Cas9 genome editing to generate a chromosomal mutation, *acn-1(am314)*. This allele has a four base pair insertion that is predicted to cause an early frameshift mutation and a strong loss of function (**Fig. 6C**). Homozygous *acn-1(am314)* mutant animals are not viable, consistent with previous reports that the *acn-1* gene has an essential function in larval development (Brooks et al., 2003; Frand et al., 2005; Oskouian et al., 2005; Metheetrairut et al., 2017). To evaluate an effect on aging, we analyzed the lifespan of *acn-1(am314)/acn-1(+)* heterozygous animals, which are viable. *acn-1(am314)/acn-1(+)* heterozygous mutants significantly extended mean lifespan by 15.1% compared to controls (**Fig. 6D, Table S5**). Thus, a partial reduction of *acn-1* activity caused a lifespan extension, indicating *acn-1* is necessary to accelerate adult lifespan. This result is consistent with results that *acn-1* RNAi, which also partially reduces gene activity, causes a lifespan extension (Kumar et al., 2016).

### The *acn-1* gene and captopril treatment interact with the *daf-16* and *daf-12* genes

*daf-16* encodes a forkhead transcription factor that functions downstream of *daf-2* (Larsen et al., 1995; Lin et al., 1997; Ogg et al., 1997). *daf-16(lf)* mutations cause a Daf-d phenotype and suppress both the Daf-c and lifespan-extension phenotypes caused by *daf-2(lf)* mutants (Gottlieb and Ruvkun, 1994). To investigate the interaction with the new *daf-2(am326)* mutation, we constructed a double mutant with *daf-16(mu86),* a strong loss-of-function allele, and treated these animals with captopril. Whereas captopril treatment extended lifespan and promoted dauer formation in *daf-2(am326)* single mutants, captopril failed to extend lifespan or induce dauer formation in these double mutant animals (**Table S5**). Thus, *daf-16* is necessary for the lifespan extension and Daf-c phenotype caused by *daf-2(am326)* and/or captopril. Kumar *et al*. (2016) reported that captopril and *acn-1* RNAi do not cause a lifespan extension in *daf-16(lf)* mutants, indicating that the activity of *daf-16* is necessary for these lifespan extending treatments; here, we confirm the results from captopril treatment (**Fig. 7A, Table S5**) (Kumar et al., 2016). However, captopril treatment does not induce nuclear localization of DAF-16 (Kumar et al., 2016). To explore the mechanism of DAF-16 activity, we analyzed the expression of a *daf-16* target gene, *sod-3* (Honda and Honda, 1999; Yanase et al., 2002; Landis and Murphy, 2010), using RT-qPCR in adult day 1 animals. Captopril treatment did not increase *sod-3* expression levels (**Fig. S3A**), suggesting that captopril treatment does not activate DAF-16 transcriptional activity. To determine if the effect might be age specific, we analyzed young (adult day 1) and middle-aged (adult day 5) animals using a fluorescent *sod-3p::gfp* transcriptional reporter. Captopril treatment did not significantly increase *sod-3* expression at either stage (**Fig. S3B**). Thus, captopril treatment does not cause nuclear localization of DAF-16 or the activation of the *sod-3* target gene, and yet *daf-16(lf)* mutations block the lifespan extension caused by the drug.

**Figure 7:**
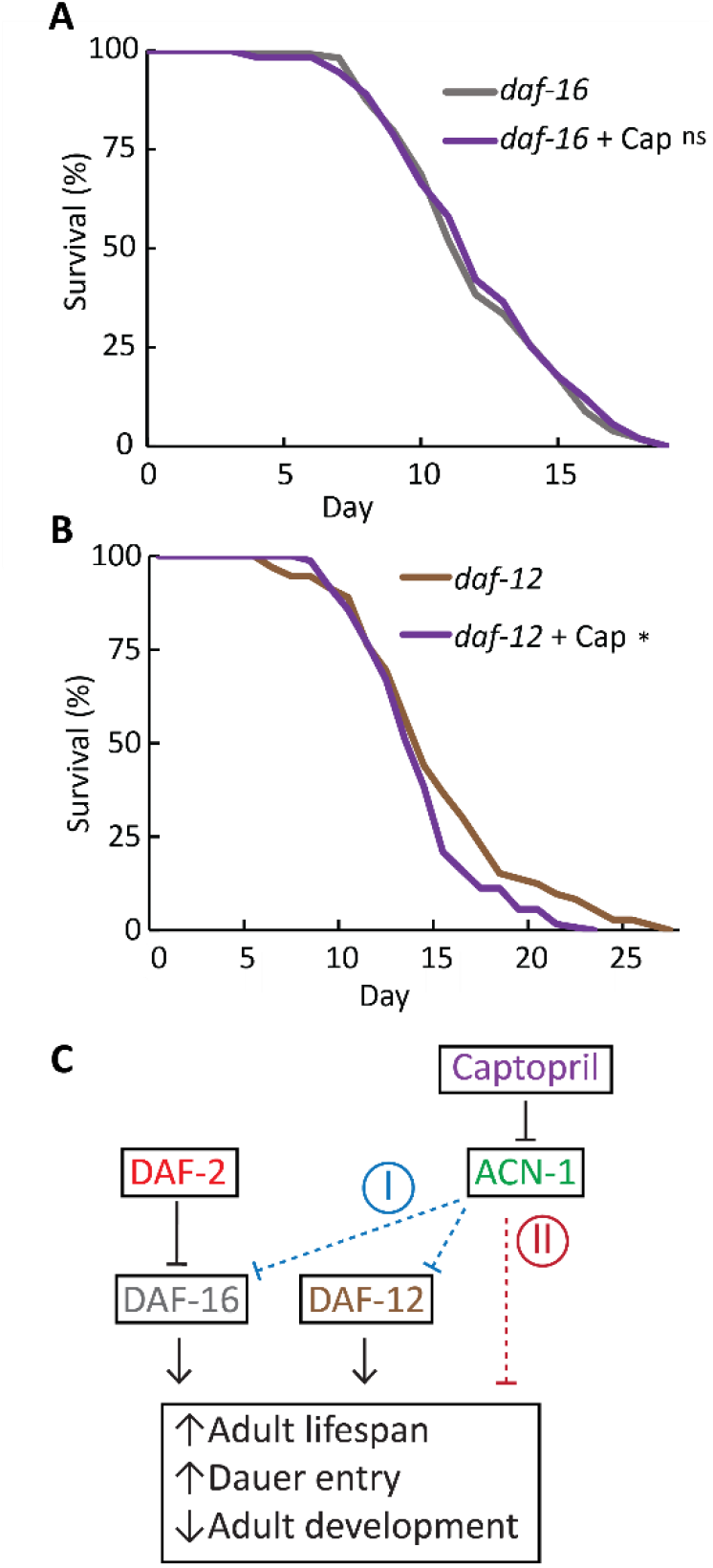
Captopril interacts with *daf-12* and *daf-16*. **(A,B)** *daf-16(mu86) and daf-12(rh61rh411)* are both loss-of-function mutations. Mutant animals were cultured with 0 or 2.5 mM captopril (+ Cap). Values indicate the fraction of the starting population that remains alive. Captopril treatment caused a small yet significant reduction in mean lifespan in *daf-12* mutants and no significant change in *daf-16* mutants. ns, nonsignificant, p≥0.05; *, p<0.05 (see **Table S5** for statistics). **(C)** A model for captopril/ACN-1-mediated control of aging and dauer formation. The DAF-2 receptor-tyrosine kinase inhibits the DAF-16 transcription factor (bar), thereby reducing lifespan, inhibiting dauer formation, and promoting larval development into reproductive adulthood. The DAF-12 transcription factor acts parallel to this pathway. ACN-1 functions to reduce adult lifespan, inhibit dauer formation, and promote larval development into reproductive adulthood; captopril functions by inhibiting ACN-1. We propose two models for ACN-1. I, blue: ACN-1 inhibits the activity of both the DAF-16 and DAF-12 transcription factors, reducing their ability to increase lifespan, promote dauer formation, and inhibit reproductive development. II, red: ACN-1 acts parallel to DAF-16 and DAF-12 to affect these phenotypes independently.

The *daf-12* gene encodes a nuclear hormone receptor that functions in parallel to DAF-16 to promote dauer formation and transcription of longevity-inducing genes (Gottlieb and Ruvkun, 1994; Ludewig et al., 2004; Shostak et al., 2004; Fisher and Lithgow, 2006). *daf-12(lf)* mutations reduce lifespan and cause a Daf-d phenotype, similar to *daf-16(lf)* mutations (Riddle et al., 1981; Vowels and Thomas, 1992; Thomas et al., 1993; Antebi et al., 1998, 2000; Ludewig et al., 2004; Fisher and Lithgow, 2006). To investigate the interaction with captopril, we treated *daf-12(rh61rh411)* mutants with the drug. Captopril treatment did not extend the lifespan of *daf-12(rh61rh411)* mutants, indicating *daf-12* is necessary for captopril-induced lifespan extension (**Fig. 7B, Table S5**). Thus, the activity of both *daf-16* and *daf-12* are necessary for the lifespan extension caused by captopril, linking two genes involved in dauer formation with the lifespan extension effects of captopril.

## Discussion

### The *C. elegans* DAF-2(A261V) mutant protein corresponds to the human INSR(A119V) mutant protein that causes Donohue syndrome

An important avenue of research in the field of aging is the identification and characterization of genes which control aging. Insulin signaling controls aging across species – reducing the activity of IIS pathway components in *C. elegans, Drosophila,* and mice extends lifespan and delays age-related degeneration (Kenyon et al., 1993; Clancy et al., 2001; Tatar et al., 2001; Taguchi and White, 2008; Bartke, 2008; Kenyon, 2010; Kannan and Fridell, 2013). In humans, this relationship is more complex, as reduction of insulin signaling is most often associated with diabetes and reduced lifespan. However, humans with exceptional longevity often display reduced insulin resistance (Bonafè et al., 2003), and altered insulin signaling in the brain has been associated with age-related neurodegenerative diseases such as Alzheimer’s Disease (Watson and Craft, 2003; Cole and Frautschy, 2007), pointing to a complex role of insulin signaling in human aging. The discovery and characterization of mutations affecting IIS pathway components represents an important avenue for furthering our understanding of aging.

In addition to aging, this pathway also regulates entry into the dauer state. Using an unbiased forward genetic screen, we identified a novel *daf-2* mutant and used it to characterize a previously unobserved interaction between captopril treatment, *acn-1* inhibition, and the dauer state. Much work has been done to categorize the numerous *daf-2* mutants based on their common phenotypes (Gems et al., 1998; Patel et al., 2008); these include extended lifespan, ectopic dauer formation, thermotolerance, and a reduction in the percentage of unfertilized oocytes compared to wild type. Subsets of *daf-2* mutants display additional phenotypes, including larval arrest, gonad abnormalities, reduced motility, and reduced fecundity. Our results suggest *daf-2(am326)* is a class II allele, based on constitutive dauer larvae formation at 22 and 25°C, extended lifespan, thermotolerance, and reduced brood size. Although *daf-2(am326)* mutants display a reduced brood size, these animals are relatively healthy in other respects. We did not observe a delay in development, reduced motility, larval arrest, gonad abnormalities, or elevated matricidal hatching. Thus, *daf-2(am326)* lacks many negative pleiotropies common to other *daf-2(lf)* mutants, and represents a useful reagent for future research on aging and insulin signaling.

The *daf-2(am326)* mutant phenotype results from an alanine to valine substitution at amino acid position 261. This residue is located in the extracellular L1 ligand binding domain of the DAF-2 protein, where a variety of ligands bind to and activate the kinase activity of DAF-2 (Kimura et al., 1997; Pierce et al., 2001; Kimura et al., 2011). Several other point mutations in this region have been shown to affect DAF-2 function, including *pe1230, gv51,* and *e979*, suggesting that relatively minor changes in protein primary structure in this region can result in large changes to protein function (Lewis and Hodgkin, 1977; Patel et al., 2008; Ohno et al., 2014; Bulger et al., 2017). This alanine residue is conserved in the human insulin receptor (INSR) at amino acid position 119 (Longo et al., 2002). Interestingly, the precise molecular lesion observed in *daf-2(am326)* has been associated with human patients that suffer from Donohue syndrome, a disease characterized by growth restriction, morphological abnormalities, reduction of glucose homeostasis, and early death (<1 year of life) (Longo et al., 2002). Our results contribute two valuable insights regarding this A to V mutation that may be relevant to the human condition: (1) this mutation is recessive; (2) this mutation causes a strong loss of function in an intact animal. Consistent with these conclusions, cell culture experiments revealed that the A119V substitution severely reduces the ligand binding activity of the human INSR (Longo et al., 2002). Previous work by Krause and colleagues has established *C. elegans* as a useful model for human insulin receptoropathies, though the INSR A119V mutation has not previously been examined in an animal model (Bulger et al., 2017). Thus, *daf-2(am326)* represents a unique opportunity to model a human INSR receptor pathology in the experimentally powerful *C. elegans* system.

### Identification of *acn-1* as a regulator of dauer formation

The identification of compounds that delay aging is critical for the field of gerontology, and model organisms such as *C. elegans* are extremely useful for screening large numbers of potential compounds. While many compounds have been found to extend lifespan in *C. elegans,* only a few have been shown to extend lifespan in mammalian model organisms, and none have been identified in humans (reviewed in (Collins et al., 2006; Lucanic et al., 2013; Partridge et al., 2020)). Pharmacological and genetic inhibition of the RAAS pathway can extend lifespan in *C. elegans, Drosophila melanogaster*, and rodents (Egan et al., 2022). However, the mechanism has not yet been well defined. Given the well established function of the RAAS in mammalian blood pressure regulation, an appealing explanation is that RAAS inhibition extends lifespan by lowering blood pressure, thereby reducing mortality related to hypertension. However, ACE inhibitors extend lifespan even in normotensive rodents (Santos et al., 2009). Furthermore, this hypothesis fails to explain the effect in *Drosophila* or *C. elegans*, animals which lack closed circulatory systems, suggesting that this is not a general model. It appears that the ACE protein evolved prior to the evolution of the closed circulatory system, implying that ACE has an ancestral function unrelated to blood pressure regulation (Simões-Costa et al., 2005; Burggren and Reiber, 2007). The observation that ACE inhibition extends lifespan in multiple species suggests that this ancestral function may control aging. Our results indicate that ACE modulates insulin/IGF-1 signaling pathways to control aging and development.

Previous studies indicate *acn-1* as a critical regulator of molting, development, and aging (Brooks et al., 2003; Kumar et al., 2016; Metheetrairut et al., 2017), and here we demonstrate a previously uncharacterized activity: control of dauer entry. Our results are consistent with two models (**Fig. 7C**). In **Model I**, ACN-1 acts upstream in a linear genetic pathway to inhibit DAF-16 and DAF-12; the mechanism of regulation is likely indirect. Inhibition of ACN-1 by captopril or RNAi promotes the activity of DAF-16 and DAF-12, thereby increasing lifespan and dauer formation. In **Model II**, ACN-1 functions in a parallel genetic pathway to DAF-16 and DAF-12 to reduce lifespan and promote reproductive development. While our results are consistent with either model, it is notable that *acn-1* RNAi and/or captopril treatment did not cause DAF-16 nuclear localization (Kumar et al., 2016) or activation of the *sod-3* target gene which favors Model II.

During early development, *C. elegans* must measure a spectrum of external signals (food availability, population, temperature, etc.) and integrate that input into a binary output: either dauer or non-dauer. Here, we provide evidence that ACN-1 is acting as a member of the dauer decision machinery. The dauer stage is a diapause mechanism that promotes survival during long-term stresses and resource deprivation – situations that are common in the wild. Dauer animals are able to survive in conditions that are fatal to non-dauers; however, dauers are sterile, and cannot transition back into reproductive development until conditions improve. Thus, improper entry into the dauer state under conditions which would otherwise permit reproduction represents a severe malus to the animal’s fitness. Complex regulatory mechanisms have evolved to provide accurate information on environmental conditions, allowing worms to make developmental decisions that maximize fitness outcomes. The insulin/IGF-1 signaling pathway is involved in interpreting external signals and implementing changes in gene expression. The receptor-tyrosine kinase DAF-2 interacts with a variety of extracellular substrates which, under dauer-inducing conditions, result in a series of phosphorylation events that serve to inhibit the DAF-16/FOXO transcription factor. In *daf-2(lf)* mutants, DAF-16 is activated, promoting dauer formation and the expression of stress-resistance and longevity-inducing genes, such as *sod-3* (Honda and Honda, 1999; Yanase et al., 2002; Landis and Murphy, 2010). ACN-1 may function in a similar fashion to integrate or modulate external signals into developmental decision making.

To date, few compounds have been identified that induce dauer formation in *C. elegans*. Ascarosides, a structurally diverse group of small molecule signaling pheromones, are excreted under conditions of high population density (Golden and Riddle, 1984; Ludewig and Schroeder, 2018). Dafachronic acids, a group of endogenously produced steroid hormones, bind to and remove the co-repressor from the DAF-12 nuclear hormone receptor (Ludewig et al., 2004; Motola et al., 2006). In addition to these endogenously produced compounds, the small molecule dafadine induces the dauer state by inhibiting DAF-9, a key effector of dafachronic acid synthesis and DAF-12-dependent dauer entry (Luciani et al., 2011). Captopril represents a newly discovered addition to the list of compounds that induce dauer, and our results suggest that the mechanism may be through modulation of the *daf* pathway. Captopril is the only one of these compounds that is known to be biologically active in humans, and whose effects on aging are conserved across species. Numerous genetic interventions have been discovered that ectopically induce or inhibit dauer formation, but these suffer from an inability to effectively titrate their effects. In this study, we have demonstrated the effectiveness of captopril in inducing dauer formation in a concentration-dependent manner. Captopril may prove useful as a tool for future screens of dauer-regulating genes.

Numerous compounds have been discovered that extend lifespan in *C. elegans*, but few have been further shown control to aging other organisms. In searching for compounds that delay aging in humans, the ideal candidate would be a compound which reproducibly controls aging in a broad range of species; this would drastically simplify mechanistic studies of aging. Recent reports show that captopril significantly extends lifespan in genetically heterologous mice (Strong et al., 2022). Based on this evidence and our increasing understanding of the mechanism of action of captopril in worms, we propose that captopril represents a promising candidate for future anti-aging studies in humans.

## Materials and Methods

### General methods and strains

*Caenorhabditis elegans* were cultured as described by (Brenner, 1974). Unless otherwise noted, strains were maintained at 20°C on Petri dishes containing nematode growth medium (NGM) and seeded with 200 µL *E. coli* OP50 bacteria. Nematode strains are listed below. CGC, *Caenorhabditis* Genetics Center at University of Minnesota. Strains WU1975-WU1978 were created by independently backcrossing WU1939 to wild type N2 (Bristol).

| Genotype | Source | Strain |
| --- | --- | --- |
| *C. elegans*: wild type | CGC | N2 (Bristol) |
| *C. elegans*: *daf-2(am326) III* | This study; 3X backcross | WU1939 |
| *C. elegans*: *daf-2(am326) III* | This study; 4X independent backcross | WU1975 |
| *C. elegans*: *daf-2(am326) III* | This study; 4X independent backcross | WU1976 |
| *C. elegans*: *daf-2(am326) III* | This study; 4X independent backcross | WU1977 |
| *C. elegans*: *daf-2(am326) III* | This study; 4X independent backcross | WU1978 |
| *C. elegans*: *daf-2(am326) III;daf-16(mu86) I* | This study | WU2008 |
| *C. elegans: daf-2(syb4952) III* | This study; SUNY Biotech | PHX4952 |
| *C. elegans: daf-2(e1370) III* | CGC | CB1370 |
| *C. elegans: daf-2(e1370) III;daf-16(mu86) I* | CGC | DR1309 |
| *C. elegans: daf-2(e1368) III* | CGC | DR1572 |
| *C. elegans: daf-2(e1371) III* | CGC | DR1568 |
| *C. elegans: daf-2(m41) III* | CGC | DR1564 |
| *C. elegans: daf-16(mu86) I* | CGC | CF1038 |
| *C. elegans: age-1(hx546) II* | CGC | TJ401 |
| *C. elegans: acn-1(am314) X* | This study | WU1746 |
| *C. elegans: daf-12(rh61rh411) X* | CGC | AA86 |
| *C. elegans: muls84 [(pAD76) sod-3p::GFP + rol-6(su1006)]* | CGC | CF1553 |
| *C. elegans: spe-9(hc88) I* | CGC | BA671 |

### Captopril and ethosuximide culture conditions

Captopril (*N-*[(S)-3-Mercapto-2-methylpropionyl]-L-proline, C4042, Sigma-Aldrich, Burlington, MA) was stored as a powder, dissolved in water at 75 mg/mL, and filter sterilized. Captopril-infused NGM dishes were prepared as previously described (Kumar et al., 2016): captopril was added to liquid agar at ∼50°C immediately prior to dispensing into dishes. Captopril-infused dishes were stored at 4°C for up to one month. For captopril dose-response experiments, embryos were placed onto *E. coli* seeded captopril dishes and observed after 72 hrs. Adult and larval animals were scored based on morphological characteristics using a dissecting microscope.

Ethosuximide (2-Ethyl-2-methylsuccinimide, E7138, Sigma-Aldrich, Burlington, MA) was similarly stored and prepared (based on methods previously described in (Evason et al., 2005; Collins et al., 2008)).

### Measurement of internal captopril concentration

High Performance Liquid Chromatography/Mass Spectroscopy (HPLC-MS) was performed on whole animal lysate. 20-30 wild-type adult hermaphrodites were cultured on medium containing 2.5 mM or 10 mM captopril and allowed to lay eggs for 2 hrs; the adults were removed, and the eggs were cultured at 20°C for 72 hrs. To remove captopril that is outside the animals, we washed animals off dishes using 1 mL water, pelleted animals by centrifugation, and removed the supernatant. Three successive wash steps were performed, and then the worms were incubated in 1 mL water on the benchtop for one hour to allow excess captopril in the intestinal lumen to be secreted. A sample of this liquid was retained for analysis, and three further wash steps were performed. The size of the worm pellet was estimated to be 2.5 µL based on the criteria and technique established in Evason *et al*. (2005) (Evason et al., 2005). The animals were lysed with Proteinase K (S-1000-1, EZ Bioresearch, St. Louis, MO), resuspended into a final volume of 100 µL, centrifuged at high speed to pellet debris, and the supernatant was submitted for HPLC-MS.

100 ng of deuterated phenylalanine d_8_ (Cayman Chemical, Ann Arbor, MI) was spiked in 100 µL of the extract as the internal standard for quantification. At the same time, 4-point calibration samples (0.05 ng/100 µL, 0.5 ng/100 µL, 10 ng/100 µL, and 100 ng/100 µL) containing 100 ng of phenylalanine d_8_ for absolute quantification. HPLC-MS equipped with a Shimadzu autosampler (20X), Shimadzu HPLC (20A, Columbia, MD), and an Applied Biosystem Sciex API-4500 Qtrap mass spectrometer (Foster City, CA) were utilized for analysis operating in positive ion multiple reaction monitoring (MRM) mode. The transitions of Q1/Q3 ions for Captopril and phenylalanine d_8_ detections were set at 218.1/116.1 and 174.1/128.1, respectively. HPLC solvent (A: 10 mM ammonium acetate in 7:3 water/ACN; B: 10 mM ammonium acetate in 1:1 isopropanol/methanol) gradient was 75% B to 30% in 4 min with a flow rate at 1 mL/min. A HILIC LC column (Waters Atlantis 2 x 100 mm, 3 µm, Milford, MA) was used. The Sciex data system (Analyst 1.52 v) was used for instrument control and data analysis (quantification). Each sample was injected twice to get the average data. Data were analyzed using Student’s *t-*test.

### Lifespan Determination

50-60 hermaphrodites at the L4 larval stage were removed from gravid populations cultured in standard conditions onto new dishes unless otherwise noted. Beginning one day later (“Day 1”), animals were observed for spontaneous movement; if no movement was observed, they were gently stimulated with a piece of platinum wire. If no stimulated movement was observed, animals were considered dead and removed from the dish. Worms which died by matricidal hatching, extrusion of the gonad, or desiccation on the sides of the dish were censored from that day onward. Worms were moved to new dishes daily during the reproductive period, or as needed thereafter. Measurement continued until all worms expired. Statistical analyses were performed using Kaplan-Meier survival function and log-rank (Mantel-Cox) test (Kaplan and Meier, 1958; Mantel, 1966).

### Measurement of body movement and pharyngeal pumping

20-30 hermaphrodites at the L4 stage were removed from gravid populations cultured in standard conditions. Animals were evaluated daily, beginning one day later (“Day 1”). Body movement was measured by depositing a worm on a dish and immediately counting the number of sinusoidal movements completed in a 15 s span; one “body movement” was defined as the movement of the head from its left-most position to its right-most position (or vice-versa) while the animal moved either forward or backward. Pharynx pumping was counted in 15 s increments per animal. Animals were transferred to a new dish during the reproductive period or as needed thereafter. Assays were continued until all animals had expired. Living animals that displayed no body movement or pharyngeal pumping were counted as “0” for the given phenotype, while dead animals were censored. Data were analyzed using one-way ANOVA with Tukey post-hoc HSD.

### Measurement of heat and oxidative stress resistance

To measure resistance to acute heat stress, we transferred 20-30 adult hermaphrodites to standard NGM dishes for one hour. Synchronized eggs laid during this hour were moved to new dishes, 30 per dish. The eggs were cultured at 20°C for 72 hrs, shifted to 35°C, and scored as alive or dead every 12 hrs until all expired. To measure resistance to oxidative stress, we prepared NGM dishes as above and added paraquat (*N,N’*-dimethyl-4,4’-bipyridium dichloride, 36541, Sigma-Aldrich, Burlington MA) to the molten agar to a final concentration of 40 mM. Eggs obtained as described above were cultured at 20°C for 72 hrs, and then worms were transferred to paraquat-containing medium. Animals were scored as alive or dead every 12 hrs until all expired. We observed that matricidal hatching was the most common cause of death in animals exposed to paraquat; therefore, animals that died from matricidal hatching were not censored from the data analysis. Statistical analyses were performed using Kaplan-Meier survival function and log-rank (Mantel-Cox) test (Kaplan and Meier, 1958; Mantel, 1966).

### Determination of dauer formation

30-50 embryos were picked from mixed stage populations cultured in standard conditions, placed on new dishes, and maintained at 20°C or shifted to 22, 25, or 27°C. After 2-3 days, dauer larvae were counted. The morphological criteria used to define dauer were: dark coloration, elongated and radially constricted body, lack of pharyngeal pumping, and presence of oral plug (Cassada and Russell, 1975; Riddle et al., 1981; Vowels and Thomas, 1992; Hu, 2018). Animals were observed until all had formed dauer, reached adulthood, or expired. The average percentage of dauer animals in a population was calculated by dividing the number of dauer larvae by the total number of animals. Data were analyzed using one-way ANOVA with Tukey post-hoc HSD.

### Measurement of developmental rate and fecundity

To determine the percentage of embryos that can reach adulthood within 72 hrs, we allowed wild-type hermaphrodites to lay eggs on a standard NGM dish for one hour. Approximately 30 eggs were picked onto several dishes containing 0, 4, 8, 12, or 16 mM captopril and cultured at 20°C for 72 hrs. Animals were scored as larva or adult using a dissecting microscope based on morphological criteria characteristic of late-stage larvae (size, presence of developing vulva, etc.) or adults (fully formed gonad with embryos, etc.).

To determine the time to first progeny, we placed 20-30 adults on a fresh dish for one hour. 20-30 of these synchronized eggs were picked onto individual dishes and cultured at 20°C for 65 hrs. Animals were then observed each hour to determine if they laid an egg(s). The time when unhatched embryos were first observed was defined as the time to first progeny, which is equivalent to the duration necessary to complete the life cycle.

To measure total and daily progeny production, 20-30 L4 hermaphrodites (“P_0_”) were transferred individually to fresh dishes and cultured with standard conditions. P_0_ animals were transferred to new dishes every 24 hrs; dishes containing eggs were cultured for 2-3 days to facilitate progeny identification, and the number of F_1_ progeny was counted. The experiment ended when self-fertile P_0_ hermaphrodites did not produce progeny for two consecutive days. Time to first progeny and total progeny were analyzed using Student’s *t-*test; daily progeny production was analyzed using one-way ANOVA and Tukey post-hoc HSD.

### Determination of worm length

30-40 embryos were picked onto dishes with medium containing 16 mM captopril and cultured for 72 hrs. Animals were washed into Eppendorf tubes using 1 mL M9 and collected by centrifugation. After aspirating the supernatant, three further wash steps were performed using 1 mL M9 to remove excess bacteria. The animals were pipetted onto standard NGM dishes lacking a bacterial lawn. Excess liquid was allowed to dry, and the animals were allowed to crawl away from each other. Photographs of individual worms were obtained using a Leica Microsystems IC80 HD microscope and then converted to greyscale using Fiji (ImageJ, 1.53f51). Determination of length was performed using the WormSizer plugin (Moore et al., 2013), and data were analyzed using Student’s *t-*test.

### A forward genetic screen for captopril hypersensitive mutants

To perform the chemical mutagenesis, we cultured large numbers of wild-type L4 hermaphrodites on dishes, washed the animals into 15 mL conical tubes using M9 buffer, pelleted animals by centrifugation at 2000 rpm for 3 min, and aspirated excess liquid to yield a volume of 4 mL. EMS (Ethyl methanesulfanate, M0880, Sigma-Aldrich, Burlington, MA) was added to a final concentration of 50 mM, and animals were gently agitated on a shaker for 4 hrs at 20°C. To remove the EMS, we performed four wash steps: animals were pelleted, the supernatant was aspirated, and the animals were resuspended in 10 mL M9 buffer. Animals were transferred to NGM dishes seeded with *E. coli* OP50 and allowed to recover and lay eggs. F_1_ self-progeny embryos were picked onto individual dishes; one F_2_ self-progeny from each F_1_ hermaphrodite was picked onto an individual dish and cultured until a gravid population had formed. The F_3+_ embryos were extracted using alkaline hypochlorite treatment and divided equally between dishes that were untreated, or contained 16 mM captopril, 50 µM zinc chloride (229997, Sigma-Aldrich, Burlington, MA), 10 µM TPEN (N,N,N’,N’-*tetrakis*-(2-Pyridylmethyl)ethylenediamine, 616394, Sigma-Aldrich, Burlington, MA), or 500 µM manganese (II) chloride (M8054, Sigma-Aldrich, Burlington, MA); dishes were seeded with 200 µL of 5X concentrated *E. coli* OP50 bacteria. Animals were allowed to grow for 3-5 days. If a strain failed to reach adulthood and produce progeny when cultured with 16 mM captopril, but did reach adulthood and produce progeny in all other stressful condition, then it was pursued.

### Whole-genome sequencing of *daf-2(am326)* mutants

The *am326* mutation was identified with whole-genome sequencing (WGS) using an EMS density mapping strategy (described in (Zuryn et al., 2010)). Using the temperature-dependent dauer phenotype to score *am326* mutant animals, we performed three successive outcrosses to N2, generating WU1939. Next, four parallel populations of WU1939 were established, and each was serially outcrossed to N2 four times. These strains, WU1975, WU1976, WU1977, and WU1978, each contained homozygous mutations at the *am326* locus and had been outcrossed a total of 7 times.

To collect chromosomal DNA from these strains, we cultured mixed-stage populations on 6-8 dishes. When all bacteria had been consumed, the animals were collected with 1mL M9 buffer into 15 mL conical tubes, centrifuged at 2000 rpm for 5 min, and excess liquid aspirated. Four successive wash steps were performed with M9 to remove excess bacteria, and the worm pellet was snap-frozen using liquid nitrogen. Four further freeze-thaw cycles were performed to lyse the worms, and the lysate was treated with Proteinase K in a 55°C water bath for 3 hrs. To remove excess worm debris, we centrifuged the lysate at 2000 rpm for 10 min and removed the supernatant from the pellet. Chromosomal DNA was extracted using the DNeasy Blood and Tissue Kit (69504, Qiagen, Venlo, Netherlands). 4 µg of DNA per sample were submitted to the Washington University Genome Technology Access Center (GTAC) for whole genome sequencing. Libraries were constructed using the KAPA Hyper PCR-free Kit (KR0961, KAPA Biosystems, Wilmington, MA) and sequencing was performed using a NovaSeq S4 XP (Illumina, San Diego, CA) at a coverage depth of ∼30X.

### Analysis of whole-genome sequencing data

Analysis of whole-genome sequencing data was performed using an EMS-density mapping approach described in (Zuryn et al., 2010; Zuryn and Jarriault, 2013). First, we aligned the reads to the WS220.64/ce10 *C. elegans* reference genome originally derived by the Genome Institute at Washington University (WUSTL) and the Sanger Institute. Reads were aligned using the bowtie2 read-alignment tool (Langmead and Salzberg, 2012). Analysis of differences between strains was performed using the MiModD v.0.1.9 toolkit (https://sourceforge.net/projects/mimodd/). Loci in the mutant strains that differed from the N2 strain were examined. Based on the logic that the *am326* mutation must be present in all backcrossed strains but absent in the N2 strain, we disregarded mutations present in some – but not all – backcrossed strains. We disregarded mutations present in the N2 strain (likely due to genetic drift causing differences between our lab N2 strain and the WS220.64 reference genome). This resulted in a list of 56 variants. Eleven variants, small insertions or deletions (“indels”) in repetitive regions, were disregarded as likely sequencing errors, since EMS mutagenesis typically creates base pair changes rather than indels. Each mutation was manually analyzed using the UCSC Genome Browser (UCSC Genomics Institute, Santa Cruz, CA) to determine the genomic region affected. See **Table S4** for detailed information on these variants. Multiple sequence alignment with human and mouse INSR and *Drosophila* InR was performed using COBALT (constraint-based multiple alignment tool) with *C. elegans* DAF-2 as the reference sequence (Papadopoulos and Agarwala, 2007).

### Complementation assays

Adult male *daf-2(am326)* or *daf-2(syb4952)* animals were mated to hermaphrodites of a different genotype at 20°C (**Table 1**). F_1_ embryo cross-progeny were shifted to 25°C and examined for dauer formation based on morphological characteristics, as described above.

### Time-series RNA-seq

Heat-sensitive sterile *spe-9(hc88)* mutant animals were synchronized by alkaline hypochlorite treatment and allowed to hatch overnight in S Basal medium. Arrested L1 larvae were transferred to 10 cm NGM dishes seeded with 200 µL *E. coli* OP50 at a density of ∼1000 animals per dish and cultured at 25°C. One dish was harvested every 24 hrs by washing with M9 buffer, after which worms were lysed with TRIzol reagent (15596026, Invitrogen, Waltham, MA) and repeatedly freeze/thawed in liquid nitrogen. Total RNA was obtained by phenol chloroform extraction and purified according to the RNeasy Micro kit protocol (74004, Qiagen, Venlo, Netherlands). RNA quality was assessed using a 4200 TapeStation system (Agilent, Santa Clara, CA). Library preparation and sequencing were carried out by Novogene Co., Ltd. (Beijing, China).

### RNAi

RNA interference was performed as described in (Kamath et al., 2003). *E. coli* HT115 bacteria expressing either a control plasmid (L4440) or a plasmid encoding *acn-1* dsRNA were obtained from the Ahringer library (Kamath et al., 2000). NGM dishes containing 50 µg/mL ampicillin (A-301-25, Gold Biotechnology, St. Louis, MO) and 1 mM IPTG (isopropyl-β-D-thiogalactoside, I2481-C50, Gold Biotechnology, St. Louis, MO) were poured and allowed to cool on the benchtop overnight. RNAi bacteria were grown overnight at 37°C in LB media supplemented with 50 µg/mL ampicillin; this starter culture was diluted 1:100 into LB media supplemented with 50 µg/mL ampicillin and grown for 6 hrs at 37°C. 200 µL was seeded onto the NGM RNAi dishes. Dishes were allowed to dry on the benchtop at room temperature overnight, then stored at 4°C. Embryos were transferred to RNAi dishes and cultured at 20, 22, or 25ׄ°C until they either reached adulthood or formed dauer larvae.

### CRISPR-mediated generation of the *am314* mutation in the *acn-1* locus

CRISPR-mediated modification of the *acn-1* locus was performed using the Co-CRISPR methodology described in (Arribere et al., 2014; Kocsisova et al., 2018). Our goal was to introduce a frameshift into the first exon of the endogenous *acn-1* locus, thus eliminating ACN-1 activity. The following mixture was injected into the gonads of P_0_ wild-type hermaphrodites: 50 ng/µL Cas9-expressing PDD162 (gift from Mike Nonet); 20 ng/µL *dpy-10* guide plasmid pMN3153; 500 nM *dpy-10(cn64)* ssDNA repair template AFZF827; *acn-1* guide plasmids (pANS1, 40 ng/µL each); 600 nM ssDNA repair template (pANS2). We selected F_1_ progeny displaying the *Rol* phenotype. *acn-1(am314)* WU1746 was derived using pyrosequencing to select heterozygous F_2_ progeny. Lifespan assays were performed by picking L4 hermaphrodites from this population as described above. To confirm the genotype of the animals that were analyzed, we performed pyrosequencing on the dead animals, and censored animals that did not display the *acn-1(am314)/acn-1(+)* genotype.

### Determination of *sod-3* gene expression levels by RT-qPCR

To measure relative gene expression, we collected whole-worm RNA, transcribed RNA into cDNA, and analyzed gene expression levels using real-time quantitative polymerase chain reaction (RT-qPCR). To isolate whole worm RNA, we picked approximately 400-500 unhatched embryos from dishes cultured in standard conditions onto 10 cm NGM dishes, which contained either 0 or 2.5 mM captopril and were seeded with 2 mL of *E. coli* OP50 bacteria. Worms were cultured at 20°C for 72 hrs and collected by washing four times with M9. Worms were compacted by centrifugation at 2000 rpm for 5 min, and excess liquid was removed via aspiration. Tubes containing the worms were frozen in liquid nitrogen and stored at -80°C. Worms were lysed by four successive freeze-thaw cycles in liquid nitrogen. 500 µL Trizol and 100 µL chloroform were added to the lysed worm solution. The mixture was vortexed and then centrifuged at 12,000 g for 15 min at 4°C. The clear supernatant layer was extracted and mixed with 500 µL isopropanol, vortexed, and centrifuged at 12,000 g for 15 min at 4°C. The RNA pellet was washed with 1 mL of 70% ethanol and centrifuged as before. Excess ethanol was aspirated, and the pellet was dried in a 60°C heat block. The pellet was resuspended in 50 µL nuclease-free water, and the RNA concentration was measured by UV spectroscopy (NanoDrop One Microvolume UV-Vis Spectrophotometer, ND-ONE-W, Thermo Fisher Scientific, Waltham, MA). Reverse transcription was performed using the iScript cDNA Synthesis Kit (1708891, Bio-Rad, Hercules, CA) using 1 µg RNA (50 ng/µL). RT-qPCR was performed using iTaq Universal SYBR Green Supermix (1725121, Bio-Rad, Hercules, CA) using 200 ng of cDNA. Relative fold-change was calculated using the comparative C_T_ method (Schmittgen and Livak, 2008); *sod-3* mRNA level was normalized using the *ama-1* control gene, and data were analyzed using Student’s *t-*test.

### Determination of *sod-3* expression using a fluorescent reporter strain

Transgenic animals containing the *sod-3p::gfp* construct were cultured from the egg stage in medium containing 0 or 2.5 mM captopril for 3 days (adult day 1) or 7 days (adult day 5). To perform fluorescence microscopy, we prepared 3% agar slides, added a drop of levamisol (3 mM) (PHR1798, Sigma-Aldrich, Burlington, MA) to the agar, and added ∼10 worms per slide. As necessary, animals were oriented using an eye lash. Excess levamisol was removed, and a cover slide was applied and sealed with 3% agar. Brightfield and fluorescence images of animals were acquired using a Leica DMi8 using the same settings for control and drug treated animals in each experiment. Fluorescence intensity of each worm was measured using Fiji (ImageJ, 1.53f51). The experiments were repeated at least three times independently, and data were analyzed using Student’s *t-*test.

## Funding

This work was supported by the National Institutes of Health [R01 AG02656106A1, R56 AG072169, R21 AG058037, and R01 AG057748 to KK, GM103422 and P60-DK-20579 to FH, 5T32GM007067-44 to BME], the Postdoctoral Fellow Seed of Independence Grant from the Department of Developmental Biology at Washington University School of Medicine to FP, and the Irving Boime Graduate Student Fellowship to BME.

## Acknowledgements

We thank the *Caenorhabditis* Genetics Center (CGC) at the University of Minnesota for providing nematode strains; the Washington University Genome Technology Access Center (GTAC) for whole genome sequencing support; SunyBiotech for CRISPR/Cas9 genome editing; Michael Nonet for expertise and reagents; Heidi Tissenbaum, Gary Ruvkun, Tim Schedl, Abhinav Diwan, Heather True, Zachary Pincus, Michael Nonet, and Roberta Faccio for useful discussion and expertise.

## Author Contributions

BME and KK conceived and designed the experiments. BME, FP, XA, SCW, IGA, PH, ZW, CC, AS, SK, MM, DLS, HF, and FH performed the experiments. BME, FP, AS, MM, HF, FH, and KK analyzed the data. HF and FH provided scientific reagents and analysis tools. BME and KK wrote the paper.

## Declaration of Interests

No competing interests declared.

## Abbreviations

ACE: Angiotensin-converting enzyme
ACE2: Angiotensin-converting enzyme 2
ACN-1: ACE-like non-metallopeptidase
Daf-c: Dauer constitutive
Daf-d: Dauer defective
EMS: Ethylmethanesulfanate
HID: High-temperature induction of dauer
HPLC/MS: High-performance liquid chromatography/mass spectroscopy
IIS: Insulin/IGF-1 signaling
INSR: Insulin receptor (human, mouse)
InR: Insulin receptor (*Drosophila*)
LB: Lysogeny broth
NGM: Nematode growth medium
SDS: sodium dodecyl sulfate
SOD: Superoxide dismutase
RAAS: Renin-angiotensin-aldosterone system
RNAi: RNA interference
RT-qPCR: Reverse transcription quantitative polymerase chain reaction
WGS: Whole-genome sequencing
WT: Wild type

---

**Figure S1:**
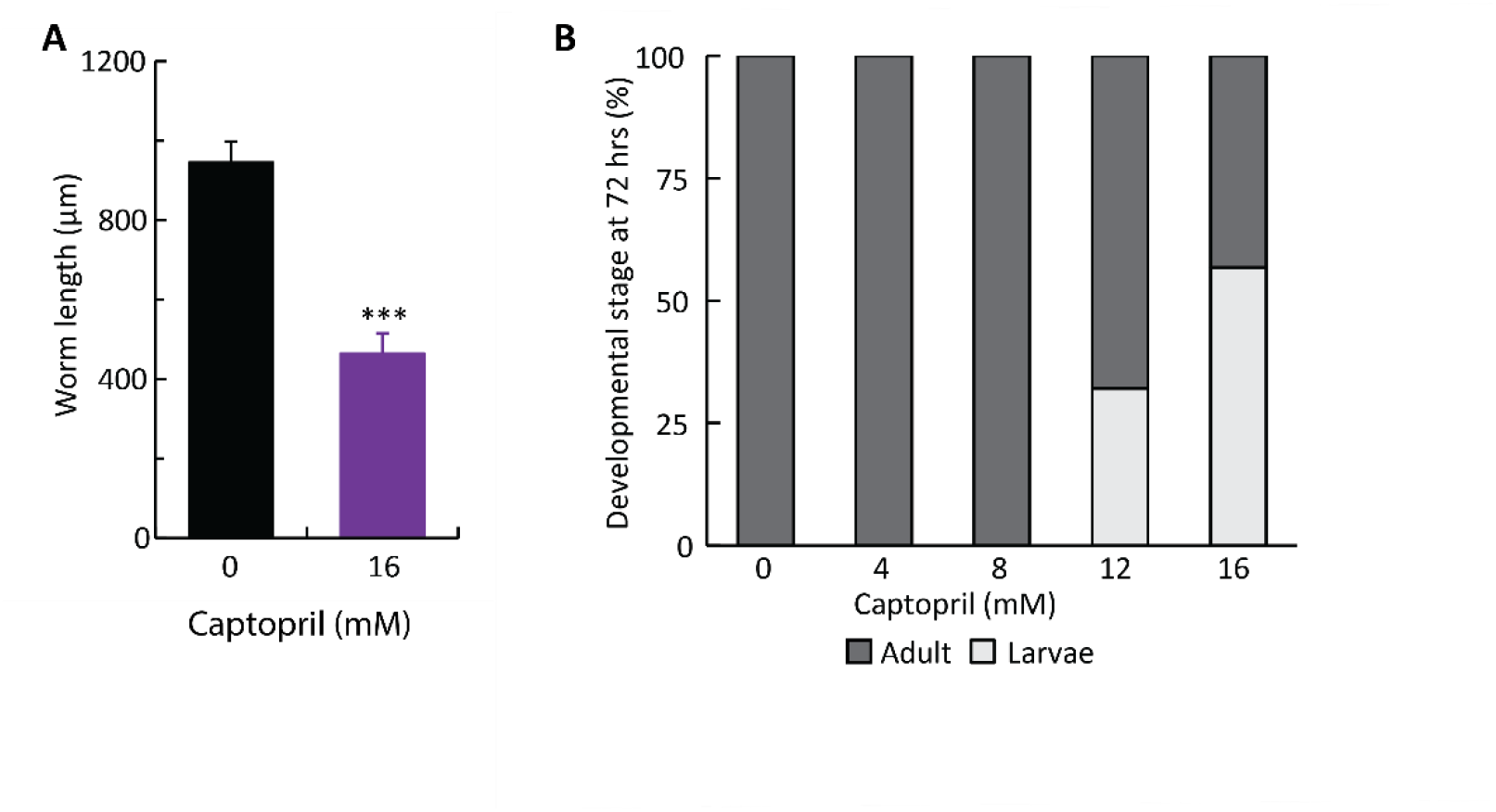
Captopril caused dose-dependent developmental delays. **(A)** Wild-type embryos were cultured on medium containing 0 (black) or 16 (purple) mM captopril for 72 hrs, and length of individuals was scored using Wormsizer (Moore et al., 2013). 16 mM captopril significantly reduced growth rate. Values are average length and s.d. n=13-32 animals, Student’s *t-*test, ***, p<0.001. **(B)** Wild-type embryos were cultured on medium containing 0, 4, 8, 12, or 16 mM captopril for 72 hrs at 20°C, and the developmental stage of each animal was classified as adult (dark gray) or larva (light gray) based on morphological criteria. Bars indicate the percentage of animals at each stage and total 100%. Captopril caused a dose-dependent reduction in the rate of development. Whereas 100% of animals matured to adults when cultured with 0, 4, or 8 mM captopril, ∼70% matured to adults with 12 mM captopril and ∼45% with 16 mM captopril. n=4 trials, N=96-119 animals.

**Figure S2:**
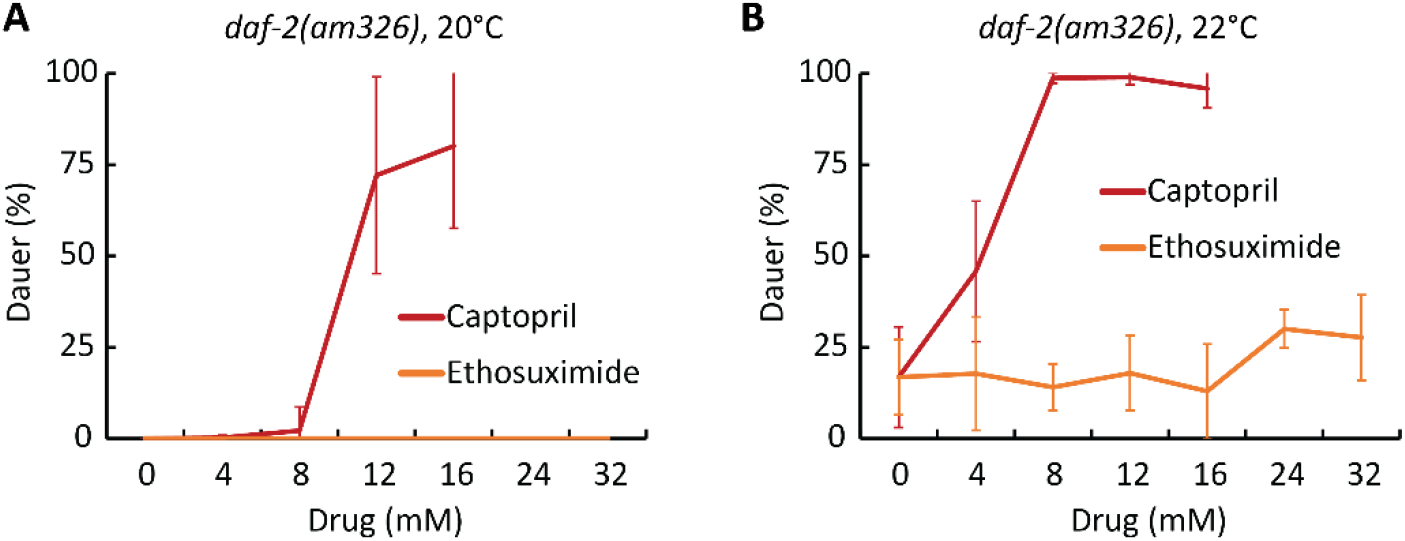
Captopril but not ethosuximide promoted dauer formation. **(A,B)** *daf-2(am326)* embryos were cultured at 20°C **(A)** or 22°C **(B)** on medium containing the indicated concentrations of captopril (red) or ethosuximide (orange). Values are the average percentage of animals in the dauer stage and s.d. (see **Table S2 and S6** for statistics). The captopril data is identical to Fig. 5A and is displayed here to facilitate comparison to the ethosuximide data. Ethosuximide is an FDA-approved anticonvulsant that extends nematode lifespan at a similar concentration to that of captopril (Collins et al., 2008; Evason et al., 2005). In contrast to captopril, ethosuximide did not promote dauer formation.

**Figure S3:**
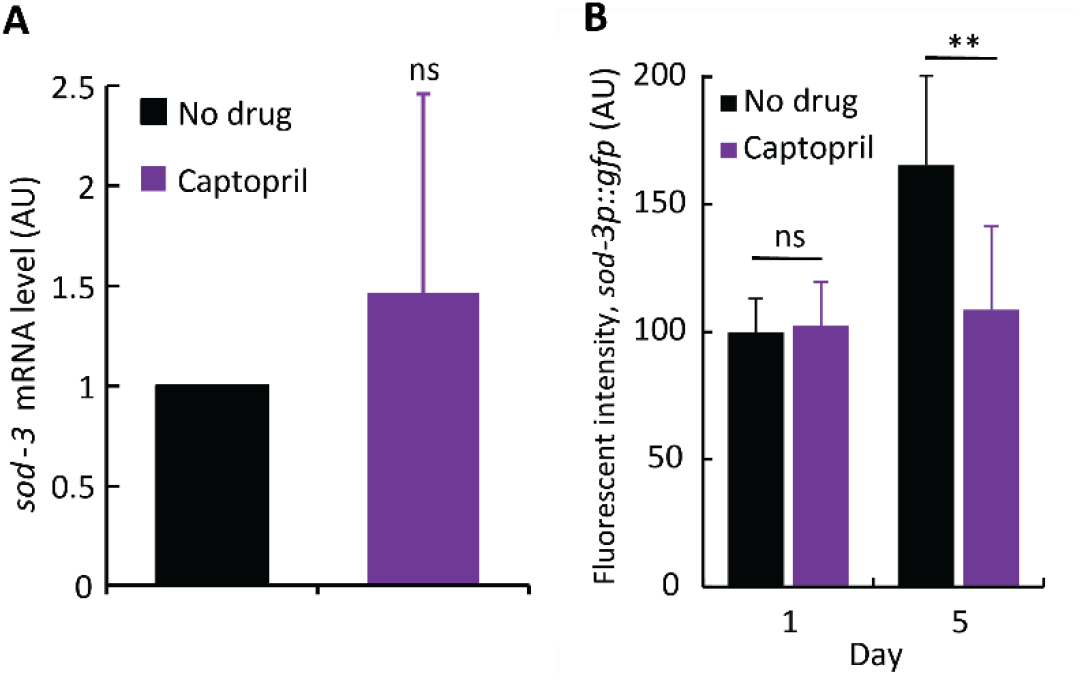
Captopril did not activate expression of *sod-3*, a DAF-16 target gene. **(A)** Wild-type embryos were cultured at 20°C on medium containing 0 mM (No drug, black) or 2.5 mM captopril (purple). After three days (day 1 of adulthood), animals were harvested for total RNA, and *sod-3* mRNA levels were measured using qPCR. *ama-1* mRNA levels were used as an internal control, and the mRNA level in 0 mM was set equal to 1.0. Values are mRNA levels in arbitrary units (AU) and s.d. No significant difference was observed. N=4 biological replicates, Student’s *t-*test. **(B)** Transgenic animals that contain a *sod-3p::gfp* transcriptional reporter were cultured at 20°C on medium containing 0 mM (No drug, black) or 2.5 mM captopril (purple). Young adult (Day 1) and middle-aged adult (Day 5) animals were analyzed by fluorescence microscopy. Values are total fluorescence quantified in arbitrary units (AU) and s.d. Captopril-treated animals did not display a significant increase in fluorescent intensity. n=3 trials, N=26-38 animals, Student’s *t-*test. ns, non-significant, p≥0.05; **, p<0.01.

**Table S1:**
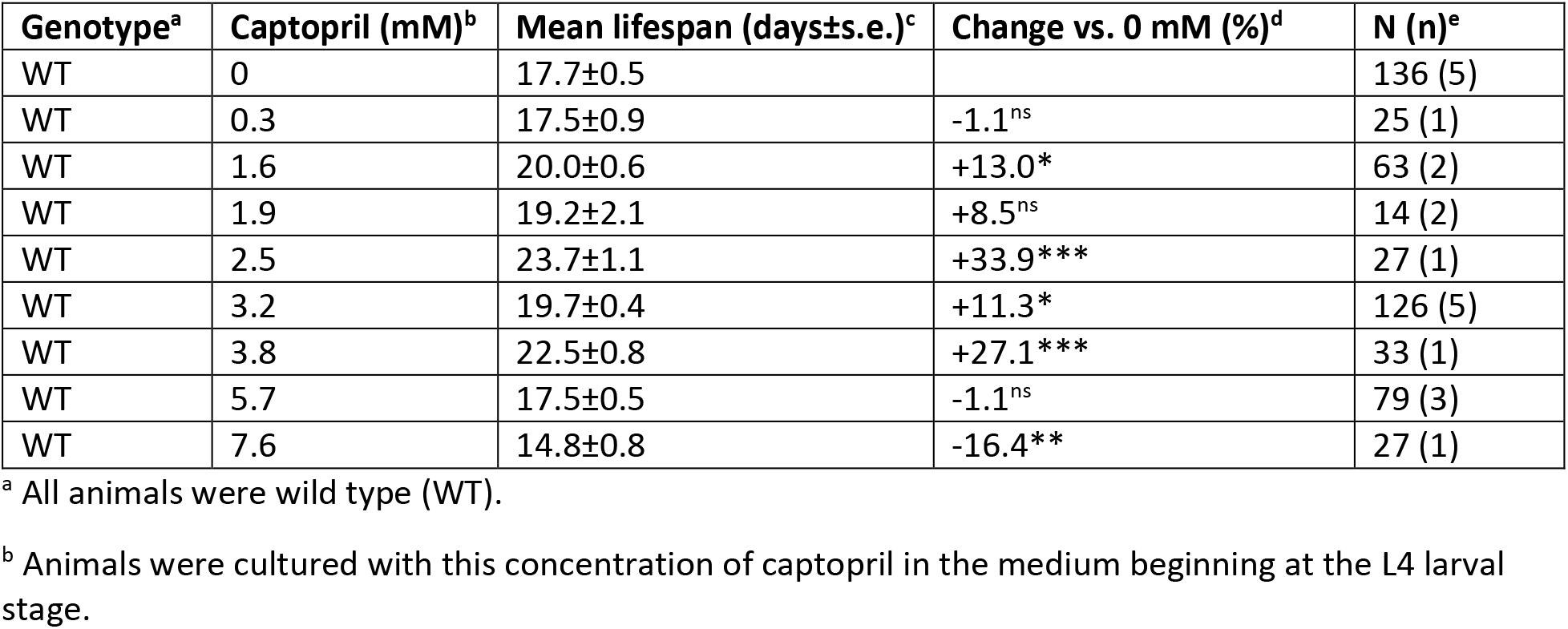
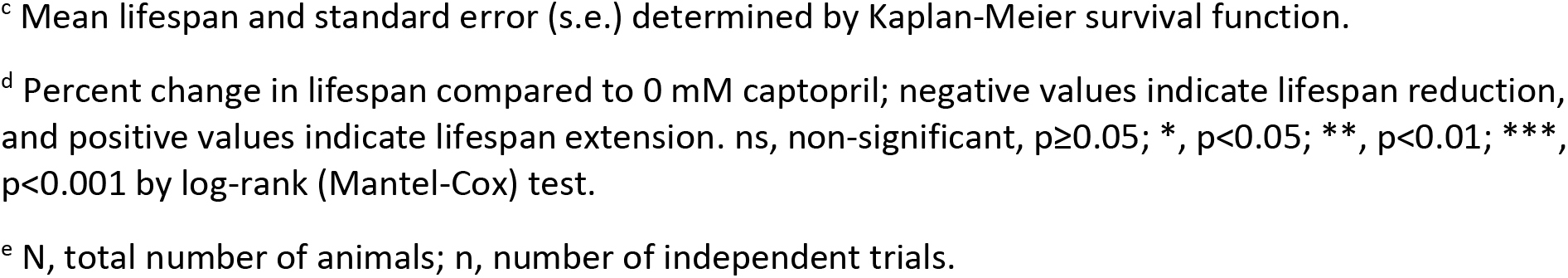
Effect of captopril on WT lifespan.

| Genotype<sup>a</sup> | Captopril (mM)<sup>b</sup> | Mean lifespan (days±s.e.)<sup>c</sup> | Change vs. 0 mM (%)<sup>d</sup> | N (n)<sup>e</sup> |
| --- | --- | --- | --- | --- |
| WT | 0 | 17.7±0.5 |  | 136 (5) |
| WT | 0.3 | 17.5±0.9 | -1.1<sup>ns</sup> | 25 (1) |
| WT | 1.6 | 20.0±0.6 | +13.0* | 63 (2) |
| WT | 1.9 | 19.2±2.1 | +8.5<sup>ns</sup> | 14 (2) |
| WT | 2.5 | 23.7±1.1 | +33.9*** | 27 (1) |
| WT | 3.2 | 19.7±0.4 | +11.3* | 126 (5) |
| WT | 3.8 | 22.5±0.8 | +27.1*** | 33 (1) |
| WT | 5.7 | 17.5±0.5 | -1.1<sup>ns</sup> | 79 (3) |
| WT | 7.6 | 14.8±0.8 | -16.4** | 27 (1) |

<sup>a</sup> All animals were wild type (WT).
<sup>b</sup> Animals were cultured with this concentration of captopril in the medium beginning at the L4 larval stage.
<sup>c</sup> Mean lifespan and standard error (s.e.) determined by Kaplan-Meier survival function.
<sup>d</sup> Percent change in lifespan compared to 0 mM captopril; negative values indicate lifespan reduction, and positive values indicate lifespan extension. ns, non-significant, $p \geq 0.05$ ; \*, $p < 0.05$ ; \*\*, $p < 0.01$ ; \*\*\*, $p < 0.001$ by log-rank (Mantel-Cox) test.
<sup>e</sup> N, total number of animals; n, number of independent trials.

**Table S2:**
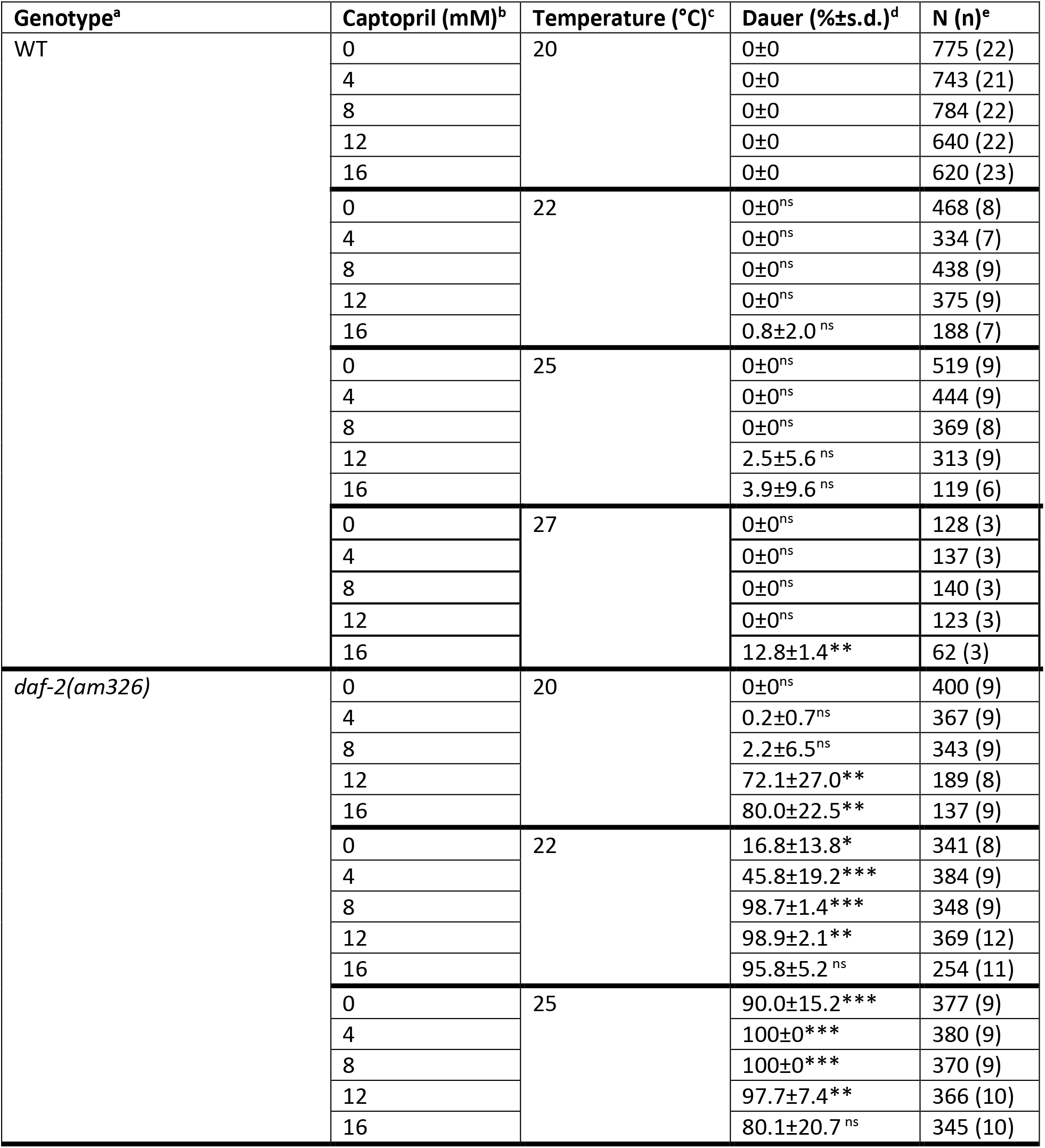
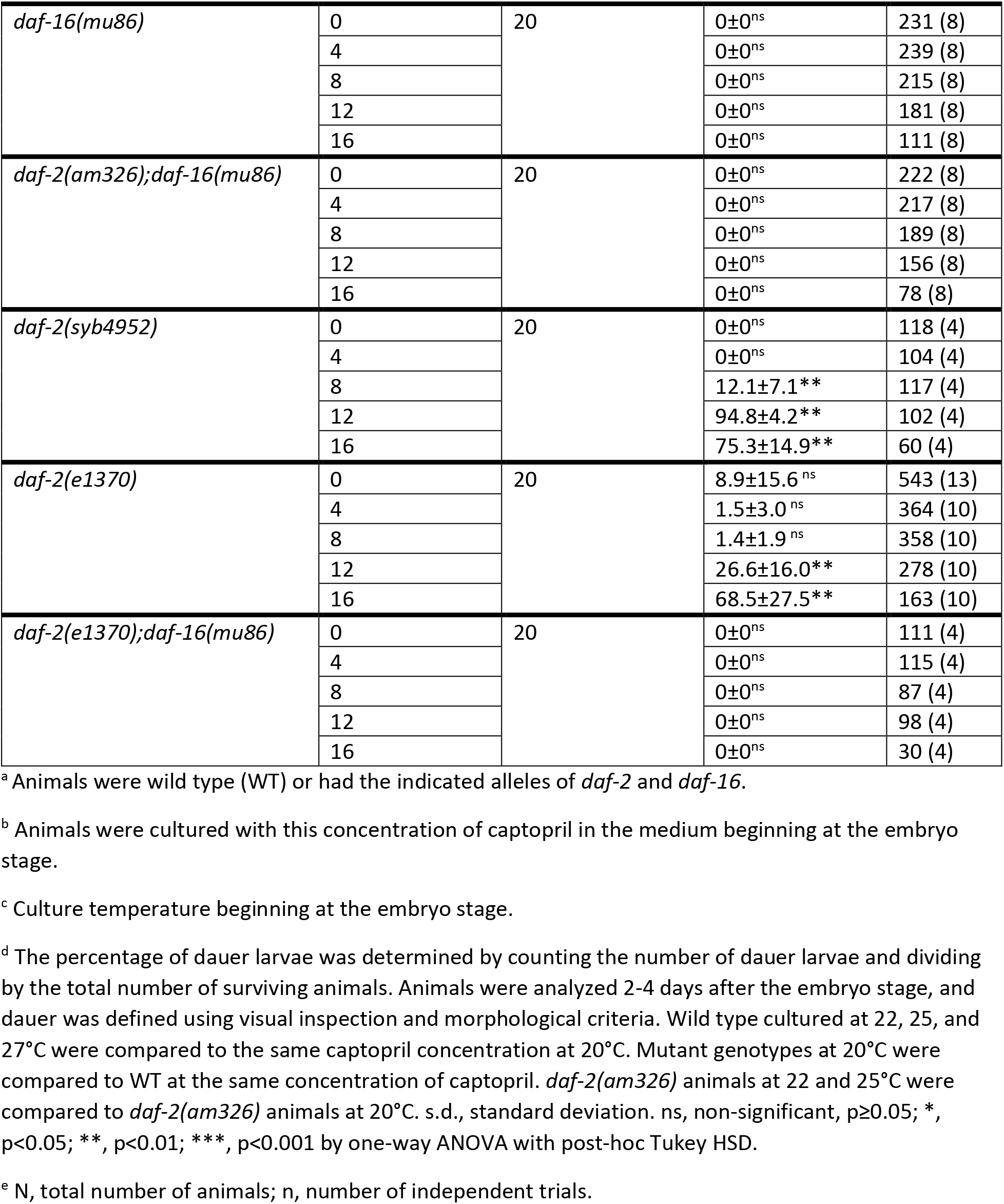
Effect of genotype, captopril, and temperature on dauer formation.

**Table S3:** Effect of captopril, oxidative stress, and heat stress on *daf-2(am326)* mutant lifespan and survival.

|  | Genotype<sup>a</sup> | Treatment<sup>b</sup> | Mean lifespan (days±s.e.)<sup>c</sup> | Survival time under stress (hrs±s.e.)<sup>d</sup> | Change (%)<sup>e</sup> | N (n)<sup>f</sup> |
| --- | --- | --- | --- | --- | --- | --- |
| 1 | WT | None | 14.5±0.2 |  |  | 365 (6) |
| 2 | WT | Captopril (2.5 mM) | 15.5±0.3 |  | +6.9*** | 219 (4) |
| 3 | *daf-2(am326)* | None | 25.3±0.5 |  | +74.5*** | 351 (6) |
| 4 | *daf-2(am326)* | Captopril (2.5 mM) | 29.5±0.8 |  | +16.6*** | 227 (4) |
| 5 | WT | Oxidation (40 mM paraquat) |  | 33.2±1.26 |  | 47 (3) |
| 6 | *daf-2(am326)* | Oxidation (40 mM paraquat) |  | 70.9±3.7 | +114*** | 57 (3) |
| 7 | WT | Heat (35°C) |  | 6±0 |  | 87 (3) |
| 8 | *daf-2(am326)* | Heat (35°C) |  | 15.7±1.0 | +162*** | 85 (3) |

<sup>a</sup> Animals were wild type (WT) or *daf-2(am326)*
<sup>b</sup> Embryos were cultured with standard medium at 20°C; beginning at the L4 larval stage, animals were transferred to standard medium at 20°C (lines 1,3), medium containing 2.5 mM captopril at 20°C (lines 2,4), medium containing 40 mM paraquat at 20°C (lines 5,6), or standard medium at 35°C (lines 7,8).
<sup>c</sup> Mean lifespan and standard error (s.e.) determined by Kaplan-Meier survival function.
<sup>d</sup> Survival time after transition of L4 animals to medium containing paraquat or high heat and standard error (s.e.) determined by Kaplan-Meier survival function.
<sup>e</sup> Percent change in lifespan or survival time. Lines 2 and 3 compared to line 1. Line 4 compared to line 3. Line 6 compared to line 5. Line 8 compared to line 7. \*\*\*, p<0.001 by log-rank (Mantel-Cox) test.
<sup>f</sup> N, total number of animals; n, number of independent trials.

**Table S4:**
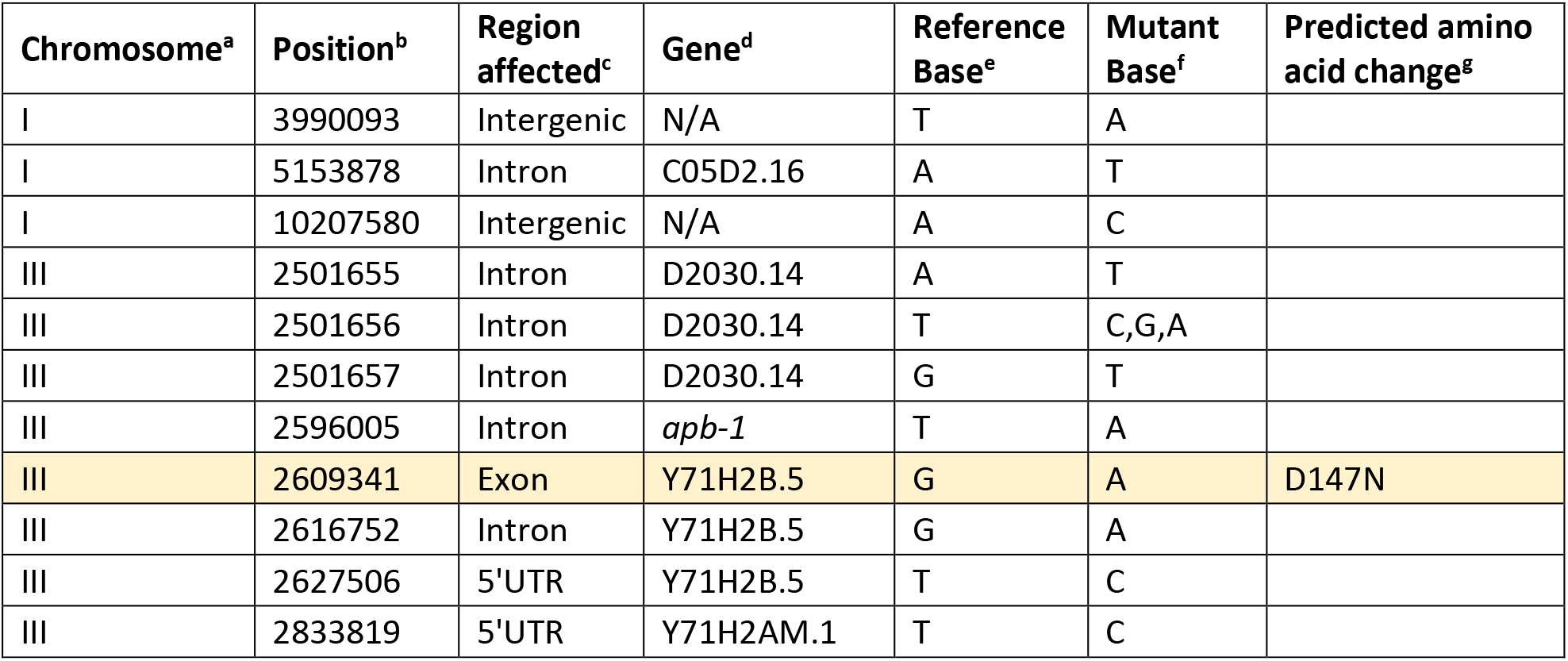
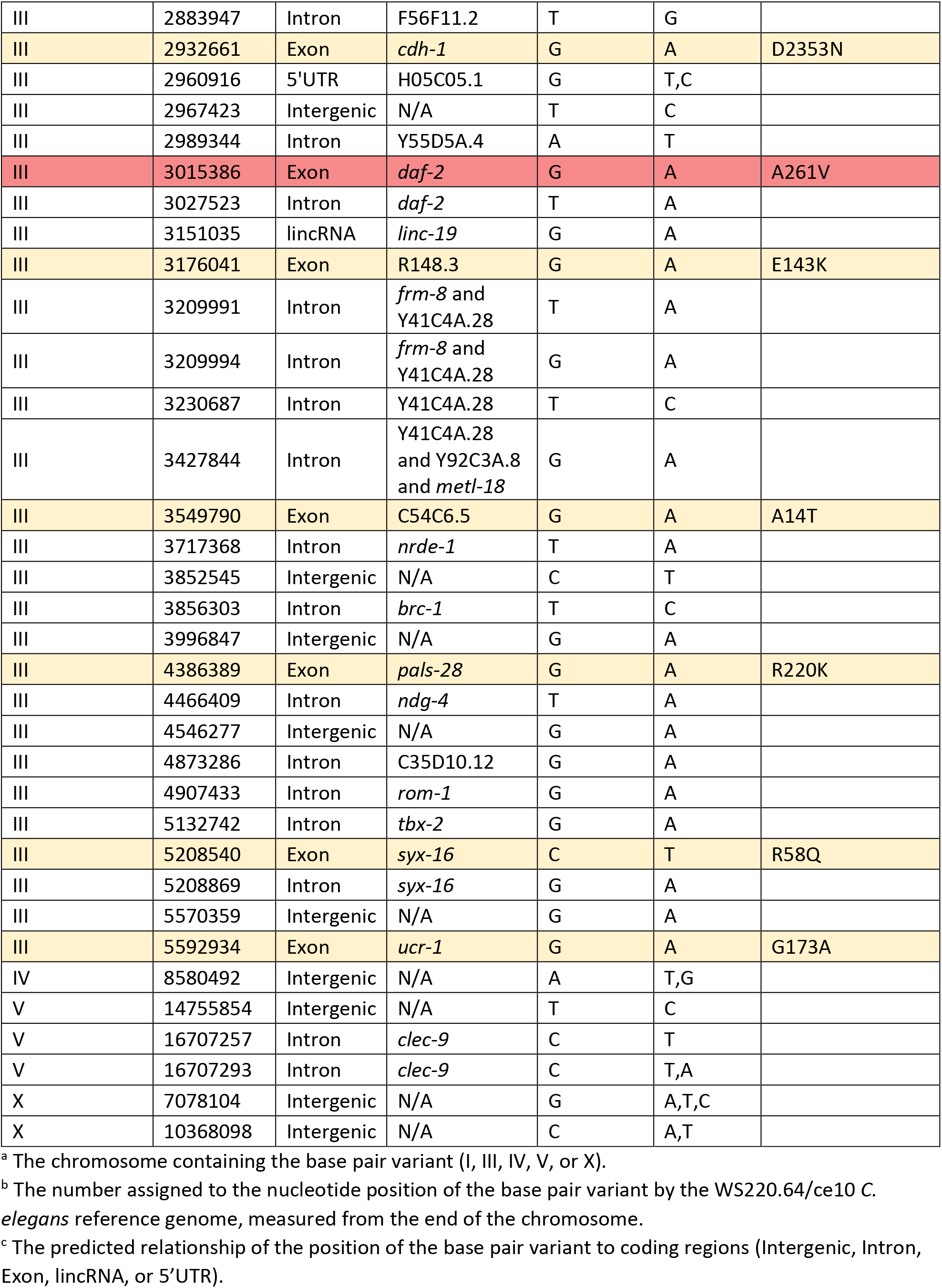
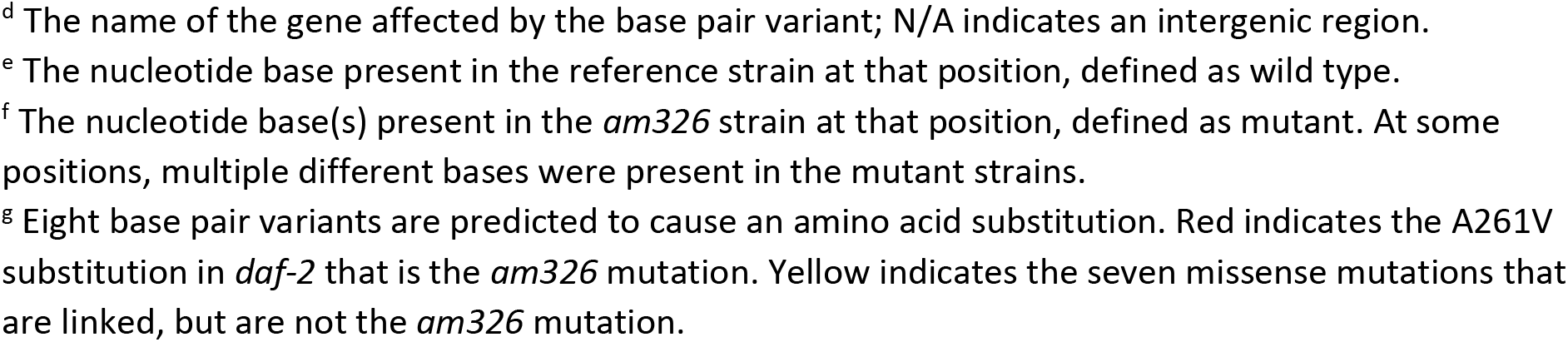
Unique base pair variants correlated with the position of *am326*.

| Chromosome<sup>a</sup> | Position<sup>b</sup> | Region affected<sup>c</sup> | Gene<sup>d</sup> | Reference Base<sup>e</sup> | Mutant Base<sup>f</sup> | Predicted amino acid change<sup>g</sup> |
| --- | --- | --- | --- | --- | --- | --- |
| I | 3990093 | Intergenic | N/A | T | A |  |
| I | 5153878 | Intron | C05D2.16 | A | T |  |
| I | 10207580 | Intergenic | N/A | A | C |  |
| III | 2501655 | Intron | D2030.14 | A | T |  |
| III | 2501656 | Intron | D2030.14 | T | C,G,A |  |
| III | 2501657 | Intron | D2030.14 | G | T |  |
| III | 2596005 | Intron | *apb-1* | T | A |  |
| III | 2609341 | Exon | Y71H2B.5 | G | A | D147N |
| III | 2616752 | Intron | Y71H2B.5 | G | A |  |
| III | 2627506 | 5'UTR | Y71H2B.5 | T | C |  |
| III | 2833819 | 5'UTR | Y71H2AM.1 | T | C |  |
| III | 2883947 | Intron | F56F11.2 | T | G |  |
| III | 2932661 | Exon | *cdh-1* | G | A | D2353N |
| III | 2960916 | 5'UTR | H05C05.1 | G | T,C |  |
| III | 2967423 | Intergenic | N/A | T | C |  |
| III | 2989344 | Intron | Y55D5A.4 | A | T |  |
| III | 3015386 | Exon | *daf-2* | G | A | A261V |
| III | 3027523 | Intron | *daf-2* | T | A |  |
| III | 3151035 | lincRNA | *linc-19* | G | A |  |
| III | 3176041 | Exon | R148.3 | G | A | E143K |
| III | 3209991 | Intron | *frm-8* and<br>Y41C4A.28 | T | A |  |
| III | 3209994 | Intron | *frm-8* and<br>Y41C4A.28 | G | A |  |
| III | 3230687 | Intron | Y41C4A.28 | T | C |  |
| III | 3427844 | Intron | Y41C4A.28<br>and Y92C3A.8<br>and *metl-18* | G | A |  |
| III | 3549790 | Exon | C54C6.5 | G | A | A14T |
| III | 3717368 | Intron | *nrde-1* | T | A |  |
| III | 3852545 | Intergenic | N/A | C | T |  |
| III | 3856303 | Intron | *brc-1* | T | C |  |
| III | 3996847 | Intergenic | N/A | G | A |  |
| III | 4386389 | Exon | *pals-28* | G | A | R220K |
| III | 4466409 | Intron | *ndg-4* | T | A |  |
| III | 4546277 | Intergenic | N/A | G | A |  |
| III | 4873286 | Intron | C35D10.12 | G | A |  |
| III | 4907433 | Intron | *rom-1* | G | A |  |
| III | 5132742 | Intron | *tbx-2* | G | A |  |
| III | 5208540 | Exon | *syx-16* | C | T | R58Q |
| III | 5208869 | Intron | *syx-16* | G | A |  |
| III | 5570359 | Intergenic | N/A | G | A |  |
| III | 5592934 | Exon | *ucr-1* | G | A | G173A |
| IV | 8580492 | Intergenic | N/A | A | T,G |  |
| V | 14755854 | Intergenic | N/A | T | C |  |
| V | 16707257 | Intron | *clec-9* | C | T |  |
| V | 16707293 | Intron | *clec-9* | C | T,A |  |
| X | 7078104 | Intergenic | N/A | G | A,T,C |  |
| X | 10368098 | Intergenic | N/A | C | A,T |  |

<sup>a</sup> The chromosome containing the base pair variant (I, III, IV, V, or X).
<sup>b</sup> The number assigned to the nucleotide position of the base pair variant by the WS220.64/ce10 *C. elegans* reference genome, measured from the end of the chromosome.
<sup>c</sup> The predicted relationship of the position of the base pair variant to coding regions (Intergenic, Intron, Exon, lincRNA, or 5'UTR).
<sup>d</sup> The name of the gene affected by the base pair variant; N/A indicates an intergenic region.
<sup>e</sup> The nucleotide base present in the reference strain at that position, defined as wild type.
<sup>f</sup> The nucleotide base(s) present in the *am326* strain at that position, defined as mutant. At some positions, multiple different bases were present in the mutant strains.
<sup>g</sup> Eight base pair variants are predicted to cause an amino acid substitution. Red indicates the A261V substitution in *daf-2* that is the *am326* mutation. Yellow indicates the seven missense mutations that are linked, but are not the *am326* mutation.

**Table S5:** Effect of captopril treatment on WT and mutant lifespan.

| Genotype<sup>a</sup> | Treatment<sup>b</sup> | Mean lifespan (days±s.e.)<sup>c</sup> | Change (%)<sup>d</sup> | N (n)<sup>e</sup> |
| --- | --- | --- | --- | --- |
| WT | None | 14.5±0.4 |  | 105 (2) |
| WT | Captopril (2.5 mM) | 17.5±0.5 | 20.6*** | 114 (2) |
| *daf-16(mu86)* | None | 12.2±0.3 | -15.9*** | 118 (2) |
| *daf-16(mu86)* | Captopril (2.5 mM) | 12.2±0.3 | 0<sup>ns</sup> | 120 (2) |
| *daf-2(am326);daf-16(mu86)* | None | 14.1±0.4 | -2.8<sup>ns</sup> | 121 (2) |
| *daf-2(am326);daf-16(mu86)* | Captopril (2.5 mM) | 14.6±0.4 | 3.5<sup>ns</sup> | 120 (2) |
| WT | None | 15.8±0.2 |  | 534 (6) |
| WT | Captopril (2.5 mM) | 17.8±0.5 | 12.7** | 185 (3) |
| *daf-2(syb4952)* | None | 30.4±0.6 | 92.4*** | 356 (5) |
| *daf-2(syb4952)* | Captopril (2.5 mM) | 34.5±0.9 | 13.5*** | 160 (3) |
| WT | None | 15.5±0.5 |  | 173 (3) |
| WT | Captopril (2.5 mM) | 17.8±0.5 | 14.8** | 185 (3) |
| *daf-12(rh61rh411)* | None | 14.8±0.4 | -4.5<sup>ns</sup> | 138 (3) |
| *daf-12(rh61rh411)* | Captopril (2.5 mM) | 13.8±0.3 | -6.8* | 169 (3) |
| WT | None | 17.2±0.6 |  | 154 (2) |
| *acn-1(am314/+)* | None | 19.8±0.7 | 15.1** | 187 (2) |

<sup>a</sup> Animals were wild type (WT) or had the indicated alleles of *daf-2*, *daf-12*, *daf-16*, or *acn-1*.
<sup>b</sup> Embryos were cultured with standard medium at 20°C; beginning at the L4 larval stage, animals were transferred to standard medium at 20°C or medium containing 2.5 mM captopril at 20°C.
<sup>c</sup> Mean lifespan and standard error (s.e.) determined by Kaplan-Meier survival function.
<sup>d</sup> Percent change in lifespan. Mutants with no drug are compared to WT with no drug. Animals treated with captopril are compared to the same genotype with no drug. Negative values indicate lifespan reduction, and positive values indicate lifespan extension. ns, non-significant, $p \geq 0.05$ ; \*, $p < 0.05$ ; \*\*, $p < 0.01$ ; \*\*\*, $p < 0.001$ by log-rank (Mantel-Cox) test.
<sup>e</sup> N, total number of animals; n, number of independent trials.

**Table S6:**
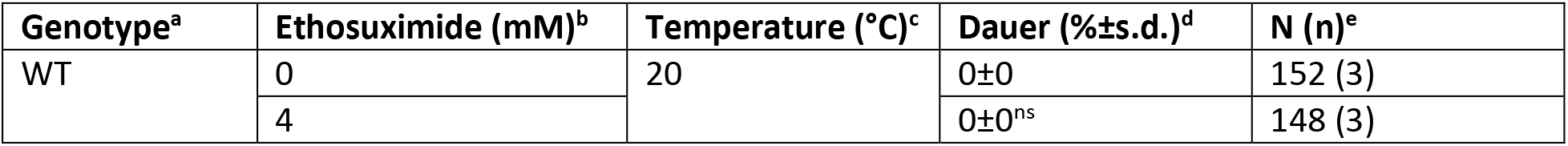
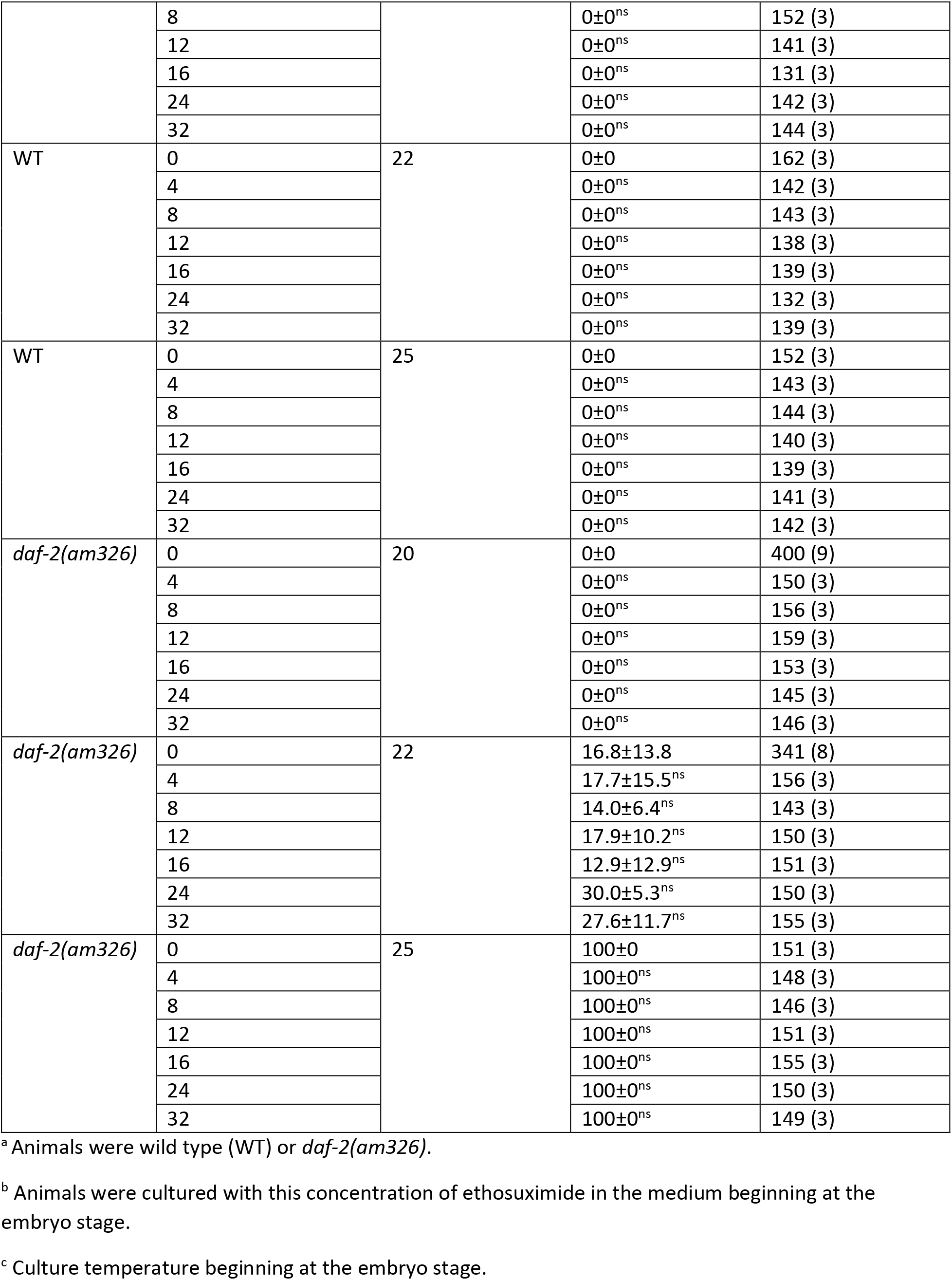
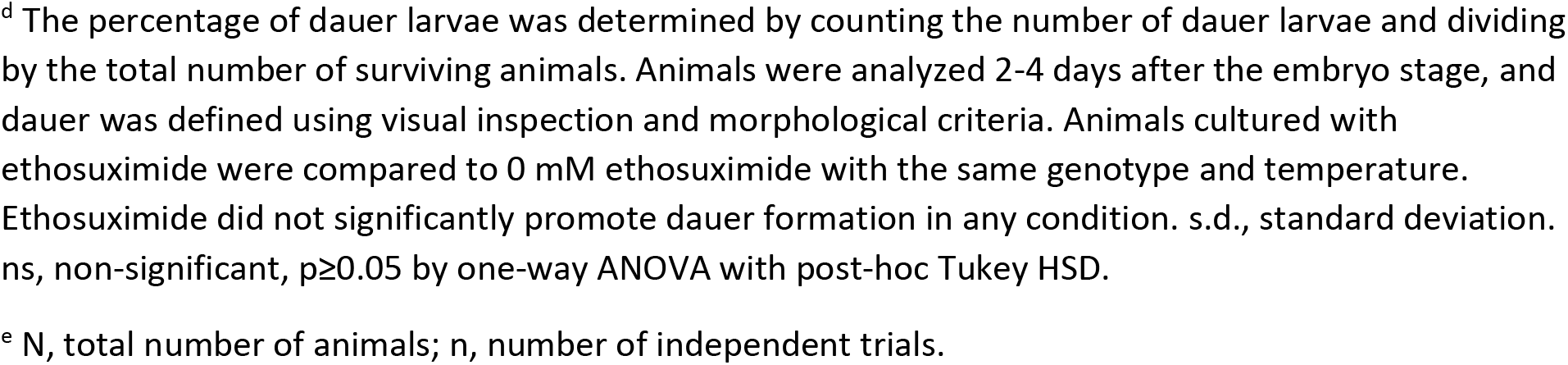
Effect of ethosuximide treatment on dauer formation.

**Table S7:** Effect of *acn-1* RNAi on dauer formation.

| Genotype<sup>a</sup> | RNAi<sup>b</sup> | Temperature (°C)<sup>c</sup> | Dauer (%±s.d.)<sup>d</sup> | N (n)<sup>e</sup> |
| --- | --- | --- | --- | --- |
| WT | Control | 20 | 0±0 | 151 (3) |
|  |  | 22 | 0±0 | 152 (3) |
|  |  | 25 | 0±0 | 152 (3) |
|  | *acn-1* | 20 | 0±0<sup>ns</sup> | 158 (3) |
|  |  | 22 | 0±0<sup>ns</sup> | 162 (3) |
|  |  | 25 | 0±0<sup>ns</sup> | 152 (3) |
| *daf-2(am326)* | Control | 20 | 0±0 | 167 (3) |
|  |  | 22 | 13.8±6.4 | 155 (3) |
|  |  | 25 | 100±0 | 155 (3) |
|  | *acn-1* | 20 | 0±0<sup>ns</sup> | 144 (3) |
|  |  | 22 | 31.5±7.4** | 149 (3) |
|  |  | 25 | 100±0<sup>ns</sup> | 154 (3) |

<sup>a</sup> Animals were wild type (WT) or *daf-2(am326)*.
<sup>b</sup> Animals were cultured with *E. coli* HT115 bacteria containing plasmid L4440 modified to encode *acn-1* dsRNA or plasmid L4440 with no insert as a control.
<sup>c</sup> Culture temperature beginning at the embryo stage.
<sup>d</sup> The percentage of dauer larvae was determined by counting the number of dauer larvae and dividing by the total number of surviving animals. Animals were analyzed 2-4 days after the embryo stage, and dauer was defined using visual inspection and morphological criteria. Values for *acn-1* RNAi were compared to control at the same temperature and genotype. s.d., standard deviation. ns, non-significant, $p \geq 0.05$ ; \*\*, $p < 0.01$ by one-way ANOVA with post-hoc Tukey HSD
<sup>e</sup> N, total number of animals; n, number of independent trials

## Notes

### Competing Interest Statement

The authors have declared no competing interest.

